# Adiponectin exerts sex-dependent effects on lipid, amino acid, and glucose metabolism during caloric restriction

**DOI:** 10.1101/2025.01.20.633882

**Authors:** Yoshiko M Ikushima, Kuan-Chan Chen, Richard J. Sulston, Domenico Mattiucci, Eleanor J. Brain, Stefanie A Fung Xin Zi, Karla J. Suchacki, Benjamin J. Thomas, Andrea Lovdel, Matthew Bennett, Hiroshi Kobayashi, Phillip D. Whitfield, Keiyo Takubo, Andrew H. Baker, Nicholas M. Morton, Robert K. Semple, William P. Cawthorn

## Abstract

Adiponectin is the most abundant hormone in the circulation. Plasma adiponectin decreases in obesity but increases in leanness, including during caloric restriction (CR) in animals and humans. In obesity, adiponectin deficiency promotes cardiometabolic dysfunction. In contrast, the roles of adiponectin in CR, when it is at its highest, are largely unknown. To address this, we studied global adiponectin knockout (KO) in male and female mice fed either *ad libitum* (AL) or a 30% CR diet from 9-13 weeks of age. We show that adiponectin KO did not alter CR effects on body mass, body composition, or energy expenditure. However, KO unexpectedly decreased blood glucose levels during CR, both on fasting and following an oral glucose challenge. This is opposite to the effects of adiponectin deficiency in the context of *ad libitum* diet (AL) or obesity, and occurred without changes in insulin secretion or sensitivity. Moreover, adiponectin KO augmented CR-induced increases in plasma fatty acids in both sexes and, in males only, impaired systemic triglyceride clearance under CR. Indirect calorimetry further revealed that adiponectin KO alters the shifts between carbohydrate and lipid utilisation that occur during transitions between fed and fasted states. To determine potential molecular mechanisms, we investigated effects of adiponectin KO on the liver, a major adiponectin target that plays key roles entraining metabolism to nutritional state. Hepatic transcriptomics revealed that, in both sexes, adiponectin KO upregulates sterol and fatty acid synthesis genes under AL while increasing amino acid catabolic genes during CR. Together, our findings suggest that adiponectin tunes glucose, lipid, and amino acid metabolism during CR, in whole or in part through effects on the liver. The widely reported functions of adiponectin in pathological states, including obesity and insulin resistance, thus differ sharply from its roles during CR, with marked sexual dimorphism apparent for many of these functions.

## 1. INTRODUCTION

Adiponectin is a multimeric secreted protein that is among the most abundant in the plasma, with typical circulating concentrations of 1-20 mg/L^1,2^. Although it is expressed almost exclusively by adipocytes, plasma adiponectin correlates inversely with adiposity^1,2^. Low plasma adiponectin in humans is also a biomarker for increased cardiometabolic risk^1,2^; hence, intensive research over the past three decades has focused on a putative mediating role for low adiponectin in adverse cardiometabolic outcomes. Much of the direct support for this comes from murine models. In mice, adiponectin deficiency usually exacerbates obesity-induced glucose intolerance, insulin resistance, and ectopic lipid accumulation in organs such as liver, pancreas and skeletal muscle (SkM)^2^. Conversely, interventions that increase adiponectin generally protect against obesity-associated metabolic dysfunction^1,2^.

The focus on adiponectin’s role in cardiometabolic disease is understandable given the biomedical imperative. However, it seems implausible that any such role explains the apparently strong selective pressure that has led to adiponectin’s high expression and phylogenetic conservation. Adiponectin is dispensable for survival under conventional animal house conditions^1,2^, but other clues to its fundamental evolutionary role may lie in the conditions under which it is most highly expressed. These include, most prominently, chronic caloric restriction (CR), defined by sustained but non-lethal decrease in calorie intake^3,4^ - a key selective pressure over evolutionary time^3^. The ability to adapt to CR by decreasing energy expenditure, thus conserving resources, is likely to have been crucial for survival in the face of privation^3–5^.

The need to minimise negative energy balance makes the increase in adiponectin production during CR all the more remarkable given that adiponectin is already highly abundant. This circumstantially implies an important role for adiponectin in adaptation to CR. However, only three studies to date have directly sought evidence for this^6–8^. They suggest roles for adiponectin in ischaemic protection and bone loss, but did not examine metabolic adaptations. Other pertinent findings are that, during fasting, mice with transgenic suppression or deletion of adiponectin exhibit greater weight and fat loss than control mice^9,10^; but these mice were not subjected to sustained CR. Thus, whether adiponectin directly influences CR’s metabolic impact remains unknown.

Importantly, investigating the role of adiponectin, if any, in adaptation to CR is motivated by more than evolutionary curiosity: CR extends healthspan in species ranging from yeast to humans^4^ and is implemented in various ways in weight loss and healthy living regimens. Understanding the molecular mediators of the beneficial effects of CR may offer opportunities to mimic these pharmacologically.

Herein, we investigated the role of adiponectin in CR by investigating male and female adiponectin KO mice and their wild-type (WT) littermates during *ad libitum* (AL) and CR feeding. Our study reveals that adiponectin has distinct, unexpected, and sometimes sexually dimorphic effects on lipid, amino acid, and glucose metabolism under AL and CR. Our findings have implications for understanding adiponectin’s evolutionary function and its contribution to CR’s therapeutic benefits.

## 2. RESULTS

### 2.1. Plasma adiponectin increases during CR

We first tested if our CR protocol causes hyperadiponectinaemia in WT mice. Compared to AL controls, 30% CR increased plasma adiponectin by ∼50% in females and ∼70% in males from weeks 3-6 of CR (Figure 1A). Females had higher adiponectin than males, irrespective of diet; however, the overall CR effect was similar between the sexes.

**Figure 1.**
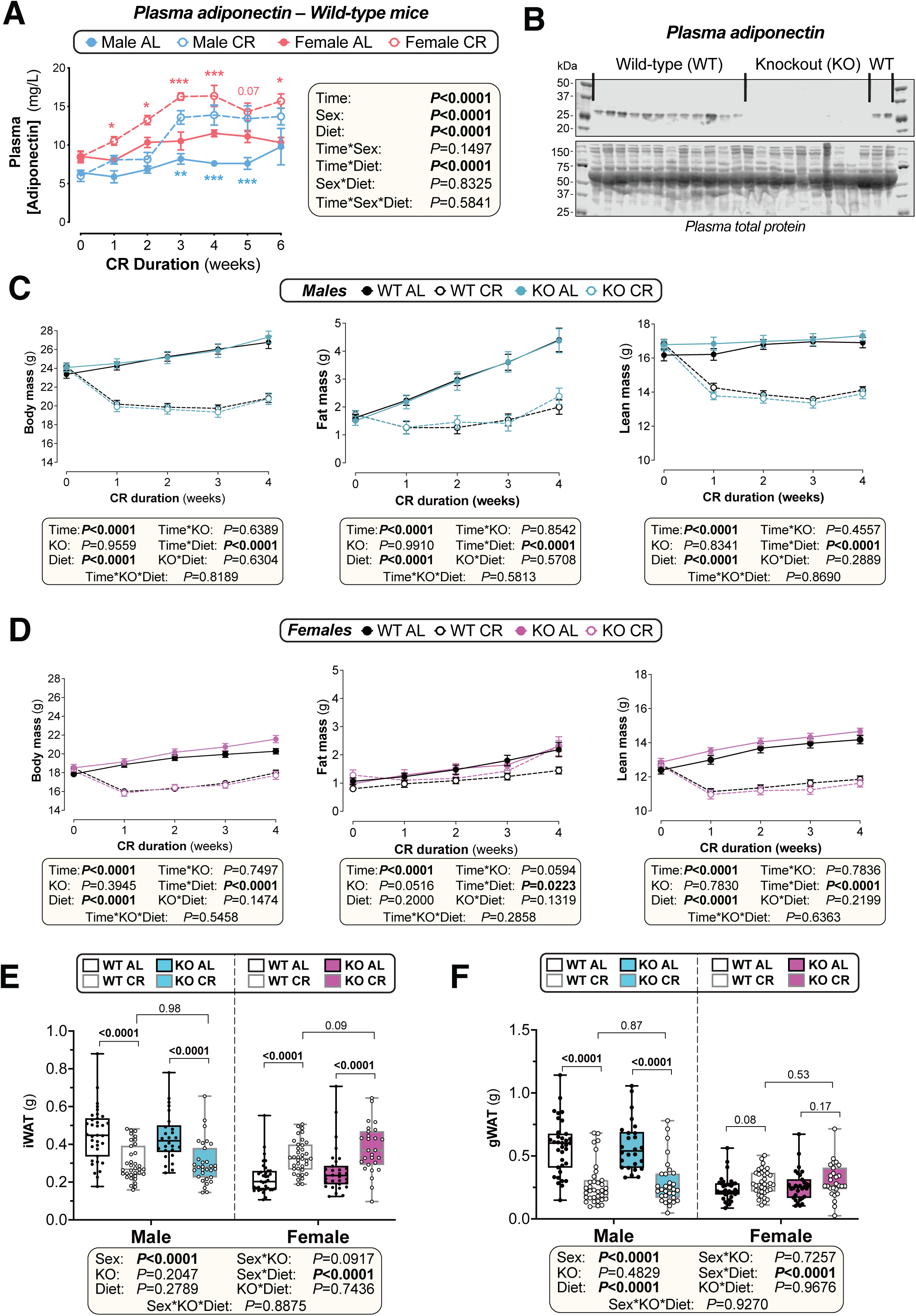
Plasma adiponectin increases during CR but adiponectin KO does not alter CR’s effects on body mass, fat mass or lean mass in both sexes. Male and female WT and *Adipoq* KO mice were fed *ad libitum* (AL) or a 30% CR diet from 9-15 weeks of age (0-6 weeks of CR) (A) or 9-13 weeks of age (0-4 weeks of CR) (C-F). (**A)** Plasma adiponectin concentrations were determined by ELISA in tail vein blood sampled weekly from male and female WT mice. Data are mean ± SEM of 5 mice per group for weeks 0-6 plus a further 5 mice per group (CR males, AL or CR females) or 6 mice per group (AL males) for weeks 0-2. **(B)** Immunoblot of adiponectin in plasma from WT and adiponectin KO mice. Equal loading was confirmed by total protein stain. (**C-D**) Each week male (C) and female (D) mice were weighed (left panel) and TD-NMR used to determine total fat mass (middle panel) and lean mass (right panel). (**E-F**) Masses of iWAT (inguinal WAT) and gWAT (gonadal WAT) were recorded at necropsy (13 weeks of age). Data are shown as mean ± SEM (C-D) or as box- and-whisker plots (E-F) of the following numbers of mice per group: *male WT AL*, n=24 (C) or 35 (E-F); *male WT CR,* n=24 (C) or 35 (E-F); *male KO AL,* n=18 (C) or 26 (E-F); *male KO CR,* n=22 (C) or 31 (E-F); *female WT AL*, n=23 (D) or 36 (E-F); *female WT CR,* n=25 (D) or 37 (E-F); *female KO AL,* n=22 (D) or 31 (E-F); *female KO CR,* n=20 (D) or 27 (E-F). Significant effects of diet, time, sex, and/or genotype, and interactions thereof, were determined by mixed-effects models (A) or 3-way ANOVA (C-F); *P values* for each variable and interaction are shown beneath each graph. In (A), 2-way ANOVA with Šidák’s multiple comparisons test was used to identify significant diet effects at each timepoint within each sex. Significant differences between AL and CR mice are indicated by * *(P*<0.05), *\*\** (*P*<0.01) or *** (*P*<0.001); blue asterisks are for males and red asterisks are for females. In (E-F), significant diet effects (within each sex and genotype) or genotype effects (within each sex and diet) were determined by 2-way ANOVA with Fisher’s LSD test; *P* values for each comparison are shown on each graph.

### 2.2. Adiponectin KO does not alter CR’s effects on body composition

We next generated adiponectin global KO mice, confirming absence of plasma adiponectin (Figure 1B, Supplementary Figure S1). WT and KO mice then underwent AL or CR feeding from 9-13 weeks of age; males and females were studied separately given our prior observation that CR exerts sexually dimorphic effects^5^. Consistent with previous findings^5^, CR decreased body mass, fat mass and lean mass in WT males whereas in females only body and lean masses decreased (Figure 1C-D). Adiponectin KO did not affect body mass, fat mass or lean mass, irrespective of sex or diet, nor, importantly, did it alter CR’s effects on these parameters (Figure 1C-D).

Decreased adiposity is critical to CR’s overall metabolic benefits^5^. As previously observed^5^, WAT in the inguinal (iWAT) and gonadal (gWAT) depots was decreased by CR in WT males but increased by CR in WT females (Figure 1E-F). Adiponectin KO did not affect iWAT or gWAT masses in either sex, irrespective of diet, nor did it alter CR’s effects on these depots. Thus, adiponectin does not influence CR’s effects on body mass, fat mass, lean mass, or regional adiposity in either sex.

### 2.3. CR-induced decreases in blood glucose are enhanced by adiponectin KO

Both CR and adiponectin increase glucose tolerance and insulin sensitivity^2,5^. Thus, we conducted oral glucose tolerance tests (OGTT) to address the hypothesis that adiponectin contributes to these CR effects. In WT mice, CR significantly lowered blood glucose levels in males and, less strongly, in females (Figure 2A-C). These data confirm our previous findings that CR has a weaker effect on glucose tolerance in females than in males^5^. However, the effects of adiponectin KO were at odds with our expectations: in males, KO *accentuated* CR-induced suppression of blood glucose across the OGTT time course, with 15-minute glucose lower in KO than WT males during CR (Figure 2A). A similar, significant KO by Diet interaction also occurred in females, despite the CR effect being weaker than in males (Figure 2B). Across both sexes, adiponectin KO enhanced CR-induced decreases in blood glucose area under the curve on OGTT (tAUC) and fasting glucose, suggesting impaired gluconeogenesis and/or increased glucose utilisation in adiponectin KO under CR (Figures 2C-D). This unexpected effect occurred without discernible alteration of either insulin secretion (Figure 2E-F) or insulin sensitivity, as assessed by the Homeostatic Model Assessment for Insulin Resistance (HOMA-IR) and insulin tolerance tests (Supplementary Figures S2A-C).

**Figure 2.**
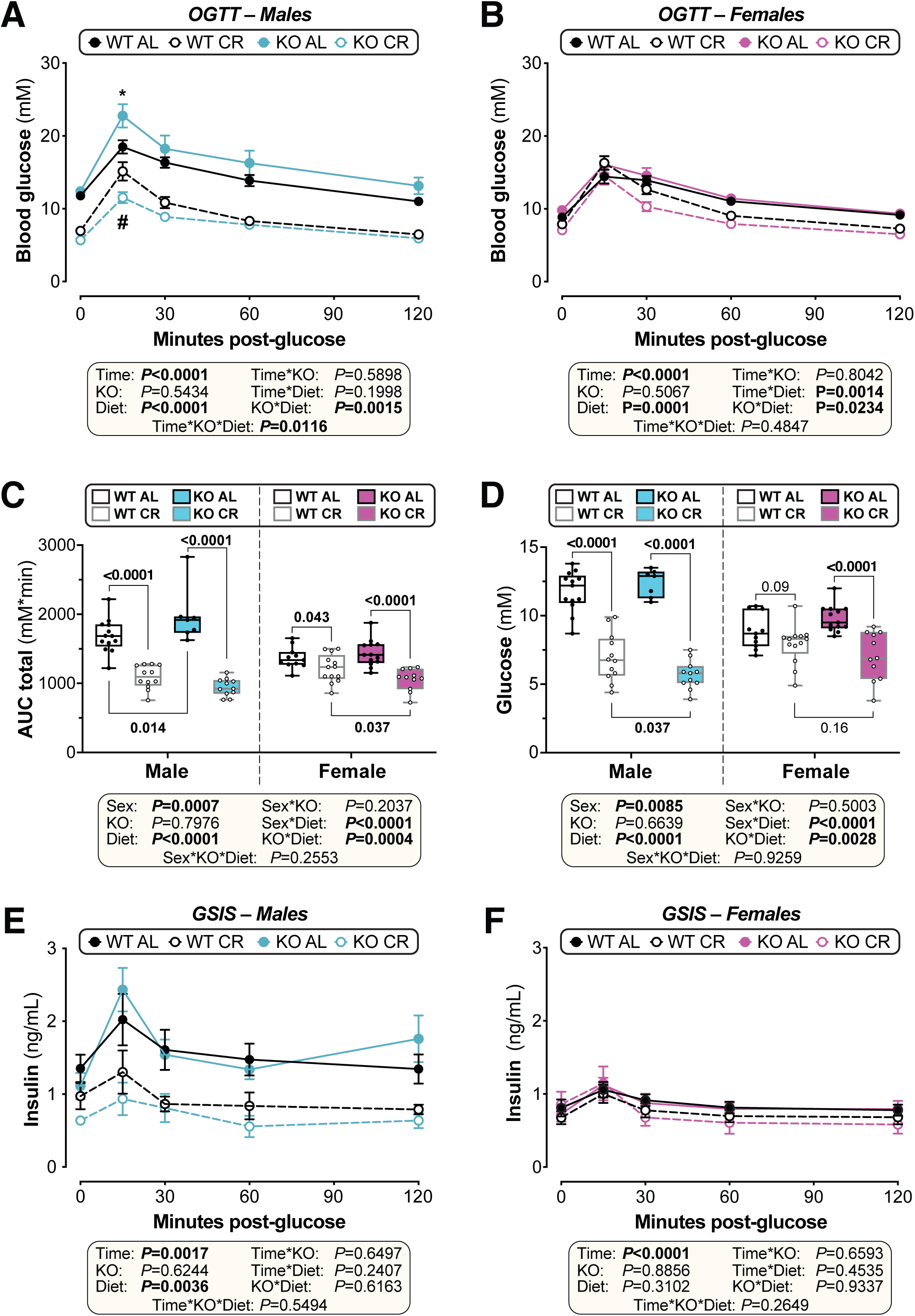
CR-induced decreases in blood glucose are enhanced by adiponectin KO in both sexes. Male and female WT and *Adipoq* KO mice were fed AL or CR as described for Figure 1. At 12.5 weeks of age (3.5 weeks of AL or CR diet) mice underwent an oral glucose tolerance test (OGTT). **(A-B)** Blood glucose readings during the OGTT for male (A) and female (B) mice. **(C)** Area under the curve (AUC) during the OGTT was determined relative to 0 mmol/L (total AUC: tAUC) for each mouse. **(D)** Fasting blood glucose concentrations at time 0 of the OGTT. **(E-F)** Glucose-stimulated insulin secretion in males (E) and females (F) during OGTT was assessed using an insulin ELISA. Data are shown as mean ± SEM (A-B, E-F) or as box-and-whisker plots (C-D) of the following numbers of mice per group: *male WT AL*, n= 13 (A,C-D) or 11 (E); *male WT CR,* n= 12 (A,C-D) or 14 (E); *male KO AL,* n= 7 (A,C-D) or 4 (E); *male KO CR,* n= 11 (A,C-D) or 4 (E); *female WT AL*, n= 11 (B,C-D) or 10 (F); *female WT CR,* n= 13 (B,C-D) or 12 (F); *female KO AL,* n= 13 (B,C-D) or 10 (F); *female KO CR,* n= 11 (B,C-D) or 7 (F). For (A-B) and (E-F), significant effects of diet, time, sex, and/or genotype, and interactions thereof, were determined by 3-way ANOVA (A-B) or mixed-effects models (E-F). For (A-B) Šidák’s multiple comparisons test was further used to assess genotype effects at 0 and 15 min; significant differences between WT and KO mice *(P*<0.05) are indicated by * or # for AL or CR diets, respectively. For (C-D), statistical analyses were as described for Figures 1E-F.

### 2.4. Adiponectin KO impairs lipid clearance and utilisation in males

Adiponectin and CR each stimulate fatty acid β-oxidation^2,3,11–14^. We next hypothesized that adiponectin contributes to increased fat oxidation during CR; that this is compromised by adiponectin KO; and that, via the Randle effect^15^, this shifts substrate oxidation towards carbohydrate utilisation, causing fasting hypoglycaemia in CR KO mice. To test this, we assessed the impact of adiponectin KO on lipid homeostasis.

Plasma non-esterified fatty acid (NEFA) concentrations increased with CR in WT males but not WT females; adiponectin KO enhanced this effect in both sexes, with CR-induced increases in plasma NEFA being greater in KO vs WT mice (Figure 3A). These data indicate that adiponectin contributes more to regulating circulating NEFA levels under CR than under AL; thus, adiponectin KO may compromise fatty acid oxidation during CR.

**Figure 3.**
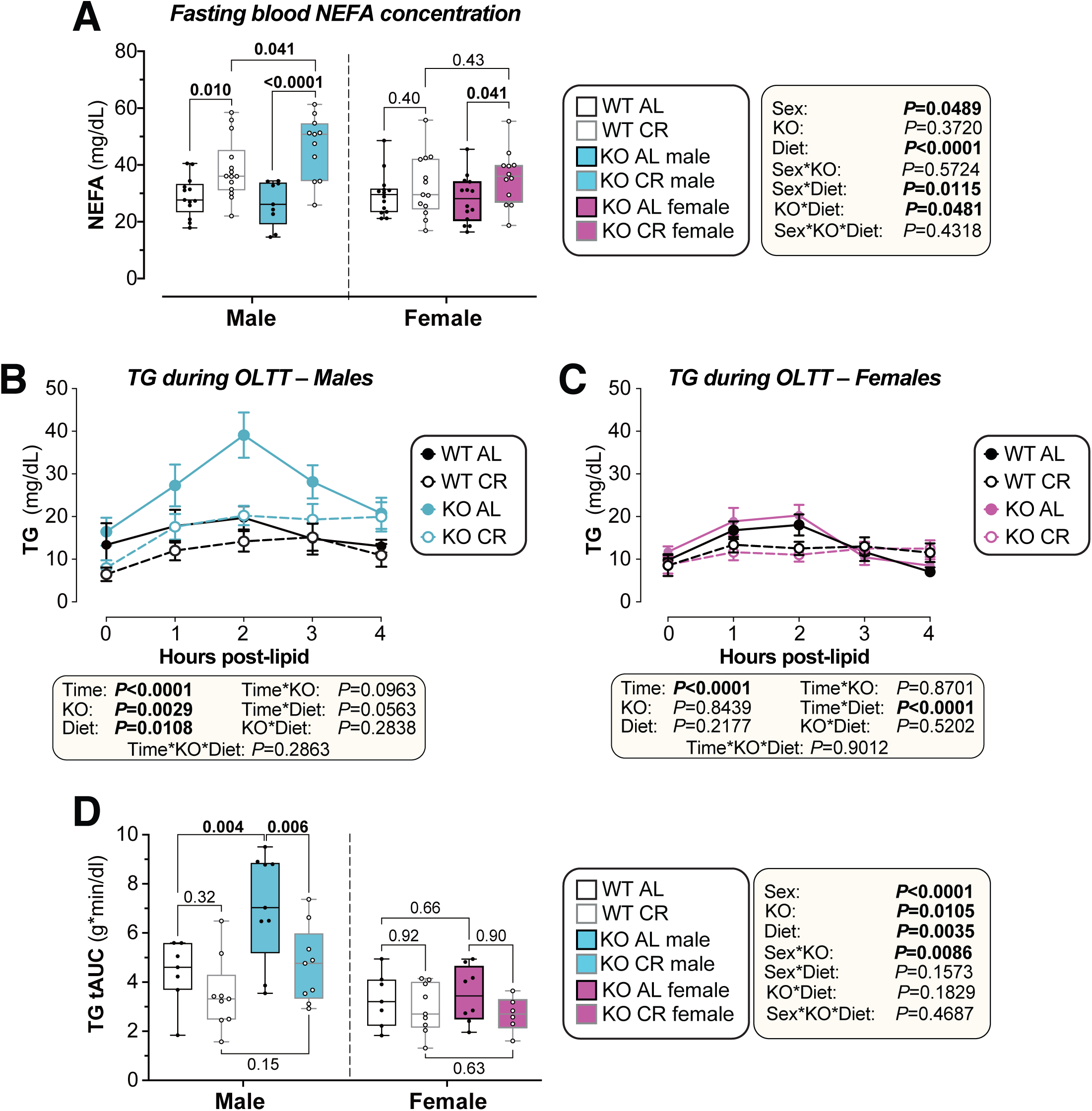
Adiponectin KO increases plasma NEFA during CR in both sexes and impairs lipid tolerance in males. Male and female WT and *Adipoq* KO mice were fed AL or CR as described for Figure 1. **(A)** Plasma NEFA measured on the day of necropsy (13 weeks of age) in mice fasted for ∼12 hours. **(B–D)** At 12.5 weeks of age (3.5 weeks of AL or CR diet) mice underwent an oral lipid tolerance test (OLTT). (B-C) Plasma triglyceride (TG) concentrations for males (B) and female (C) during the OLTT. (D) Total AUC for TG during the OLTT was determined relative to 0 mg/dL TG. Data are shown as box-and-whisker plots (A, D) or as mean ± SEM (B-C) of the following numbers of mice per group: *male WT AL*, n= 13 (A) or 7 (B, D); *male WT CR,* n= 14 (A,C-D) or 9 (B, D); *male KO AL,* n= 9 (A,C-D) or 9 (B, D); *male KO CR,* n= 11 (A,C-D) or 9 (B, D); *female WT AL*, n= 14 (A,C-D) or 7 (C-D); *female WT CR,* n= 13 (A,C-D) or 9 (C-D); *female KO AL,* n= 14 (A) or 9 (C-D); *female KO CR,* n= 12 (A) or 6 (C-D). Significant effects of diet, genotype, time and/or sex, and interactions thereof, were determined by 3-way ANOVA (A-B, D) or mixed-effects models (C). For (C-D), significant diet effects (within each sex and genotype) or genotype effects (within each sex and diet) were assessed as described for Figures 1E-F.

Previous studies show that adiponectin KO impairs lipid tolerance in ageing and other contexts^16^, but whether this occurs in CR is unknown. Therefore, we undertook oral lipid tolerance tests (OLTT). In males, plasma TG was decreased by CR but increased by adiponectin KO, indicating that KO impairs lipid clearance; this KO effect was stronger in AL vs CR males (Figure 3B). On the other hand, in females, CR prevented the increases in TG that occurred during the OLTT but adiponectin KO had no effect on lipid clearance (Figure 3C-D).

Together, these data indicate that adiponectin KO may compromise fatty acid oxidation during CR and reveal striking sexual dimorphism in the effects of adiponectin KO on lipid tolerance.

### 2.5. During CR, adiponectin KO enhances postprandial carbohydrate utilisation in males and impairs lipid oxidation in fasting females

To further interrogate the impact of adiponectin KO on substrate utilisation and metabolic homeostasis under CR, we used indirect calorimetry to assess systemic substrate utilisation and energy expenditure (EE) during week 3 of CR (Figure 4). Consistent with previous observations^5^, feeding CR mice at ∼1030 increased the respiratory exchange ratio (RER) to >1 until the beginning of the dark period, indicating postprandial fatty acid synthesis from dietary carbohydrates^17^ (Figure 4A-B). RER of CR mice reached a nadir of ∼0.7 from 0100-0900, a fasting period wherein fatty acids are the predominant fuel source. In contrast, AL mice had greater RER during the dark period and lower RER in the light period, corresponding to periods of increased and decreased feeding, respectively (Figure 4A-B). Across the full 24 h light-dark cycle, CR significantly decreased average RER in males and females (Figure 4C). In WT mice, CR also robustly suppressed 24 h energy expenditure (Figure 4D), driven largely by decreased energy expenditure during the night (Figures 4E-F).

**Figure 4.**
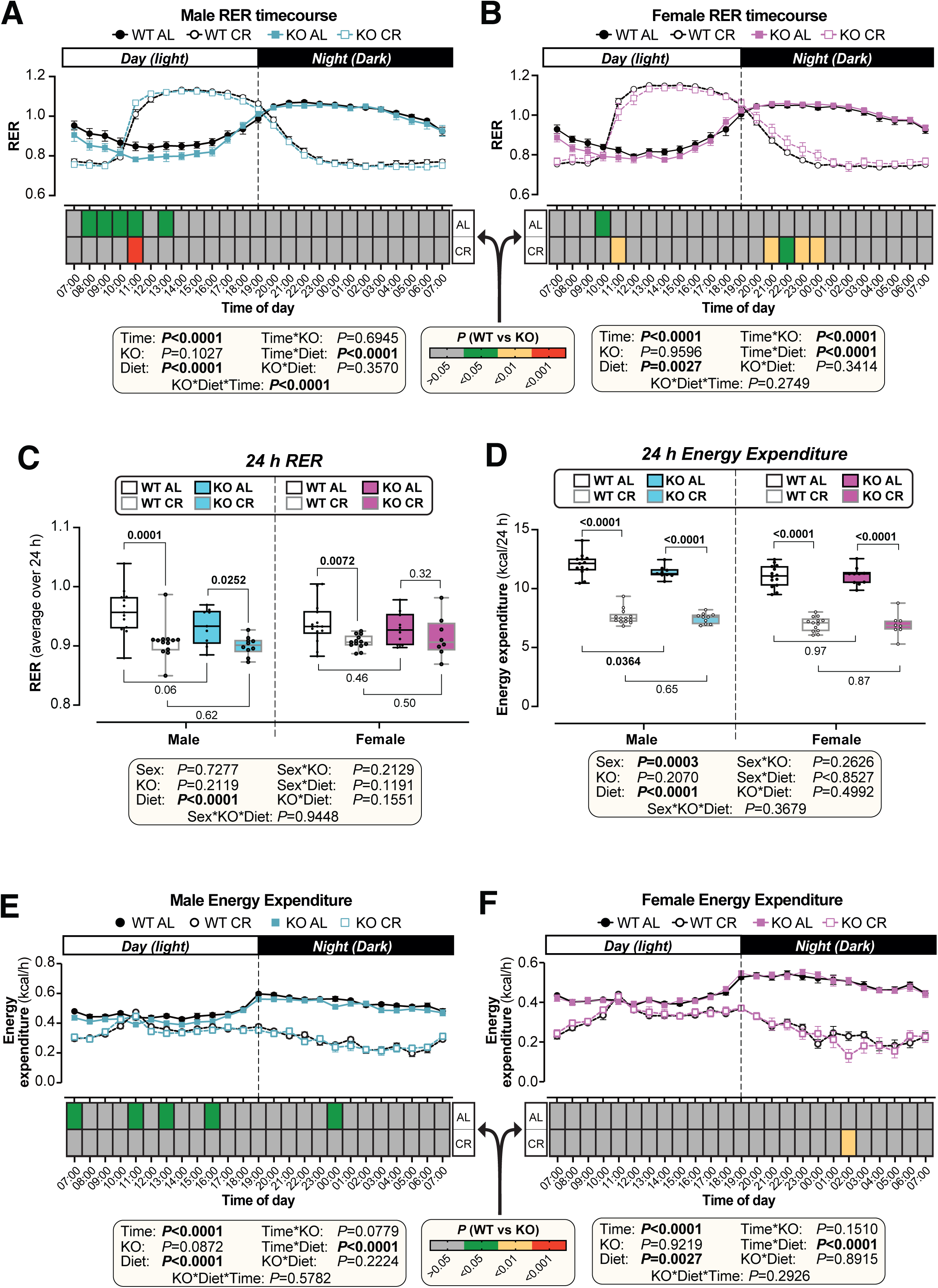
Effects of adiponectin KO and CR on RER and energy expenditure. Male and female WT and *Adipoq* KO mice were fed AL or CR as described for Figure 1. At 11.5 weeks of age (during the third week of AL or CR diet) mice were housed for 4 days in Promethion CORE System cages for indirect calorimetry. Respiratory exchange ratio (RER) and energy expenditure (EE) were recorded every minute throughout the 4 days. **(A)** Male and **(B)** Female RER per hour over the 24-hour light (Day) and dark (Night) periods, based on the average for days 2–4 of Promethion housing. **(C)** Average RER across the 24-hour light-dark cycle. **(D)** Average EE across the 24-hour light-dark cycle. **(E, F)** Male (E) and Female (F) energy expenditure per hour over the 24-hour day and night periods. Data are presented as mean ± SEM (A, B, E, F) or box-and-whisker plots (C,D) of the following numbers of mice per group: *male WT AL*, n= 13; *male WT CR,* n= 13; *male KO AL,* n= 9; *male KO CR,* n= 10; *female WT AL*, n= 14; *female WT CR,* n= 13; *female KO AL,* n= 10; *female KO CR,* n= 8. Significant effects of genotype, diet, and time or sex were determined by 3-way ANOVA. Overall *P* values from 3-way ANOVA are shown beneath each graph. For (C-D), significant genotype effects (within each sex and diet) or diet effects (within each sex and genotype) were assessed as described for Figures 1E-F. *P* values from these multiple comparisons are indicated by the colour coding for each timepoint (A, B, E, and F) or shown on the graphs (C-D).

In AL or CR mice of either sex, adiponectin KO did not alter average RER or energy expenditure over the 24 h period (Figures 4C-D) and had relatively little effect on energy expenditure at specific time points (Figures 4E-F). However, KO had diet- and time-dependent effects on RER that might reflect its effects on glucose and lipid metabolism (Figures 2-3). For example, KO decreased fasting (daytime) RER in AL males and females but increased fasting (nighttime) RER in CR females (Figures 4A and B). The latter suggests that, under CR, KO females resist lipid oxidation and/or have greater carbohydrate oxidation in the fasted state. This may explain the lower OGTT blood glucose and increased fasting NEFA in KO vs WT females during CR (Figures 2B, 3A). Moreover, during CR, adiponectin KO enhanced the postprandial increase in RER (10:00-11:00) in males but suppressed this in females (Figures 4A-B). One possibility is that, during CR, adiponectin KO enhances postprandial glucose utilisation in males. If so, this may explain why KO males, but not KO females, have lower OGTT peak glucose under CR (Figures 2A-B).

These observations demonstrate that although KO does not affect overall RER or EE in AL or CR mice of either sex, it can alter the shifts between lipid and carbohydrate utilisation that occur during transitions between fed and fasted states. Taken together, the results of our tolerance tests and indirect calorimetry suggest that adiponectin plays a role in defending against hypoglycaemia during CR by suppressing utilisation of glucose as a substrate for energy production.

### 2.6. Adiponectin KO has no effect on liver mass, triglycerides, or sphingolipid concentrations in AL or CR-fed mice

The liver plays a pivotal role in orchestrating fluxes in macronutrient metabolism on nutritional transitions, and is also a key target of adiponectin; in obese mice, recombinant adiponectin suppresses hepatic glucose production^18^, whereas adiponectin KO increases hepatic gluconeogenesis and insulin resistance^19^. This results, in part, from adiponectin decreasing hepatic steatosis and ceramide accumulation, which otherwise compromise insulin sensitivity^2^. Thus, we next investigated the hepatic effects of adiponectin KO during both ad libitum feeding and CR.

CR decreased liver mass in males but not females and adiponectin KO did not influence this, regardless of sex or diet (Supplementary Figure S3A). CR effects on liver mass occurred without changes in hepatic triacylglycerol (TG) concentrations or content, which were also largely unaltered by adiponectin KO (Supplementary Figure S3B-C); the exception was for males, in which CR decreased hepatic TG content in KO but not WT livers (Supplementary Figure 3C). Targeted lipidomics of male livers showed that CR altered hepatic ceramides, dihydroceramides (DHC), and the ceramide:DHC ratio in a manner dependent on sphingolipid chain length (Supplementary Table 1-3). Strikingly, adiponectin KO did not alter ceramides, DHCs, or the ceramide: DHC ratio in AL or CR mice, nor did it alter their modulation by CR (Supplementary Table 1-3).

### 2.7. Adiponectin KO has a major effect on lipid metabolism-related genes under AL and on amino acid catabolism-related genes under CR

To interrogate the hepatic effects of adiponectin KO in more detail, we analysed livers by bulk RNA-seq. Principal component analysis (PCA) demonstrated that sex and diet caused many more transcriptional changes than adiponectin deficiency, with WT and KO samples colocalising within each sex-diet subgroup (Figure 5A). Nevertheless, between WT and KO mice we identified 380 differentially expressed genes in AL males, 777 in CR males, 723 in AL females, and 1438 in CR females (Figure 5B-D, Supplementary Figures S4A-C, Supplementary Table 4). KO-affected genes showed minimal overlap across subgroups (Figure 5C, Supplementary Figures S4B), indicating that adiponectin’s impacts on hepatic gene expression are highly dependent on sex and nutritional state.

**Figure 5.**
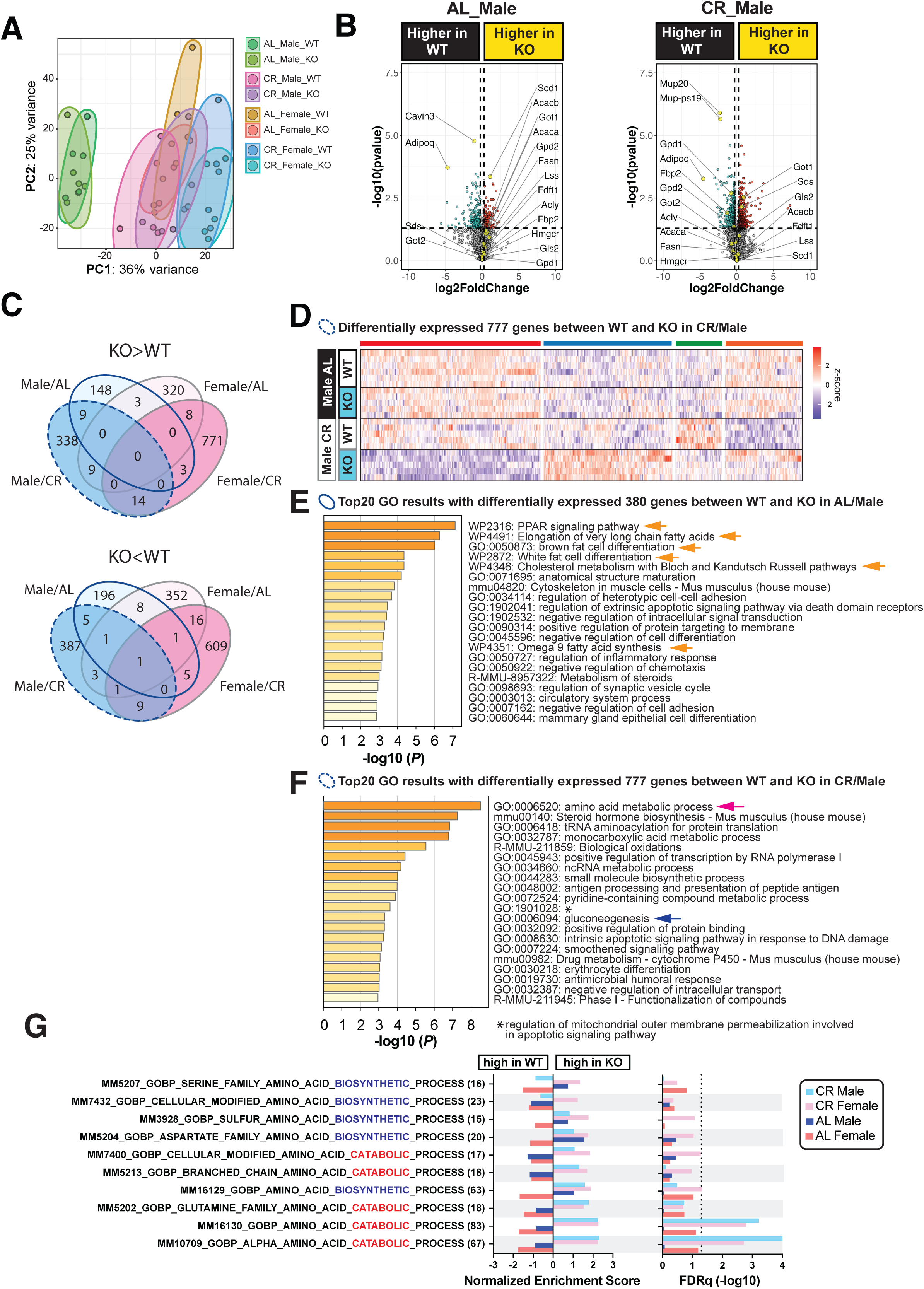
Adiponectin KO enhances hepatic expression of amino acid catabolism genes during CR in both sexes. Male and female WT and *Adipoq* KO mice were fed AL or CR as described for Figure 1. At 13 weeks of age, mice were culled and livers were collected and analysed by bulk RNA-seq. The following numbers of mice were used for each group: *male WT AL*, n= 6; *male WT CR,* n= 5; *male KO AL,* n= 5; *male KO CR,* n= 5; *female WT AL*, n= 5; *female WT CR,* n= 6; *female KO AL,* n= 6; *female KO CR,* n= 6. **(A)** Principal component analysis (PCA) for all 8 groups. **(B)** Volcano plots showing differently expressed genes between WT and KO in male diet subgroups (AL Male, CR Male). Genes with rawp value < 0.05 and absolute fold change >1.2 are shown in blue or red dots. The names of some genes of interest are shown. **(C)** Overlap of genes differentially expressed between WT and KO in each subgroup (AL Male, CR Male, AL Female, and CR Female). Genes increased in KO vs WT (rawp<0.05, fold change >1.2) are shown in the upper Venn diagram and genes decreased in KO vs WT (rawp<0.05, fold change <-1.2) are shown in the lower Venn diagram. **(D)** Clustered heat maps of the 777 genes differentially expressed (rawp <0.05, absolute fold change >1.2) between WT and KO under CR in males, the genes included in the dashed blue circles in the Venn diagrams in (C). **(E)** Results of the GO analyses conducted with the 380 genes differentially expressed between WT and KO in AL Male, the genes included in the blue circles in the Venn diagrams in (C). The top 20 clusters of GO results are shown. The yellow arrows highlight the GO term related to lipid metabolism. **(F)** Results of the GO analyses conducted with the 777 genes shown in (C) and (D). The top 20 clusters of GO results are shown. The blue arrow highlights the GO term for gluconeogenesis and the pink arrow for amino acid metabolism. **(G)** GSEA results for gene sets related to amino acid catabolism and amino acid biosynthesis in each subgroup (AL Male, AL Female, CR Male, CR Female), with comparison of WT vs KO mice. Data were extracted from GSEA results with the M5 gene set library (m5.all.v2023.2.Mm.symbols.gmt). Gene sets are arranged according to the normalised enrichment score (NES) value in CR Male. The left graph shows NESs and the right graph shows –log10 (FDRq) values. For male CR, gene set MM10709 returned an FDRq result of 0; for visualisation purposes, this is shown with a –log10 (FDRq) value of 4 in the graph. The dashed line indicates FDRq of 0.05. Genes not shown in the volcano plots in (B) because of their high rawp values or high absolute FC values are summarised in Supplementary Table 4.

Gene ontology (GO) analysis of genes differentially expressed between KO and WT revealed that, in AL males and females, adiponectin KO altered transcripts relating to lipid metabolism (Figure 5E, Figure S4D, yellow arrows). Gene set enrichment analysis (GSEA) further revealed that sterol biosynthesis-related genes were upregulated in KO under AL in both sexes (Supplementary Figure S5A and S5B). Changes in FA biosynthesis-related genes were also highly enriched in KO in both sexes under AL and in females, but not males, under CR (Supplementary Figure S5C). In contrast, during CR, amino acid metabolism-related genes were among the most significantly enriched in KO vs WT mice of both sexes (Figure 5F, Supplementary Figure S4E; pink arrows). GSEA further revealed that, among these genes, transcripts related to amino acid catabolism were particularly enriched in KO vs WT in both sexes (Figure 5G, Supplementary Figure S5D-E). Indeed, 35 genes in CR males and 39 genes in CR females showed core enrichment for the MM10709_GOBP_ALPHA_AMINO_ACID_CATABOLIC_PROCESS gene set, among which 19 genes were common between the sexes (Supplementary Figure S6A-B).

SkM is a major source of gluconeogenic amino acids during fasting^20^. Thus, we speculated that, if adiponectin KO increases hepatic amino acid catabolism, this is fuelled by amino acids from SkM. To test this, we analysed the masses of gastrocnemius and soleus muscles and myofiber cross-sectional area (CSA) of gastrocnemius muscle. Overall, muscle mass was lower in females vs males and was significantly decreased by CR (Supplementary Figure S7A-D), which also lowered gastrocnemius myofiber CSA (Supplementary Figure S8A-B). However, adiponectin KO did not affect these parameters, nor did it alter their modulation by CR (Supplementary Figures S7 and S8). Thus, adiponectin is unlikely to substantially influence SkM protein breakdown and amino acid release to the circulation under CR.

In CR males, KO also altered hepatic expression of gluconeogenesis-related genes, including highly expressed genes such as *Got1* and *Sds*, and genes with lower expression, such as *Fbp2*, *Gpd1*, and *Gpd2* (Figures 5B, 5F [gene set GO0006094, blue arrow], and Supplementary Figures 7A-B). In contrast, no gluconeogenesis-related gene sets were among the top GO results in CR females (Supplementary Figure S4E). The lower expressions of *Gpd1* and *Gpd2* in KO vs WT males during CR (Figure 5B and F, Supplementary Figure S9A-B) suggests that KO may compromise gluconeogenesis from glycerol. This could explain the lower fasting blood glucose levels in KO males under CR (Figure 2D), as glycerol is the main gluconeogenic substrate in fasting mice^21^. However, glycerol tolerance tests revealed that adiponectin KO moderately impaired gluconeogenesis in AL males only, with no KO effects in CR in either sex (Supplementary Figure S9C-D).

Together, our liver RNA-seq data demonstrate a clear contrast in the roles of adiponectin under AL and CR: in both male and female livers, adiponectin primarily targets lipid biosynthesis-related genes during AL feeding, but amino acid catabolism-related genes during CR. These hepatic effects provide further clues about the mechanisms through which adiponectin KO impacts systemic glucose and lipid metabolism and highlight that adiponectin’s functions are highly diet- and sex-specific.

## 3. DISCUSSION

Adiponectin has been studied extensively in obesity and insulin-resistant states, in which circulating adiponectin decreases. Yet adiponectin’s fundamental roles in resilience to physiological, evolutionarily relevant stressors, such as sustained negative energy balance, have been largely overlooked. Several studies report that CR in animal models or humans increases glucose tolerance and insulin sensitivity without increasing circulating adiponectin^22–26^, suggesting that hyperadiponectinaemia is not required for these metabolic benefits. We have now directly tested this by investigating the impact of adiponectin KO on the metabolic effects of CR. We show that adiponectin KO does not influence CR-induced weight loss, changes in body composition, suppression of energy expenditure, or decreases in average 24hr RER. However, unlike in obesity, during CR adiponectin KO unexpectedly decreases blood glucose during fasting and following an oral glucose load. Adiponectin KO also augments CR-induced increases in plasma fatty acids in both sexes and, in males, significantly reduces lipid clearance; the latter KO effect is stronger in AL vs CR males. These KO effects on systemic glucose and lipid metabolism are supported by our indirect calorimetry data, which suggest that adiponectin KO alters the shifts between carbohydrate and lipid utilisation that occur during transitions between fed and fasted states. Finally, our bulk liver transcriptomic analyses identify diet-dependent effects: in both sexes, adiponectin KO predominantly alters hepatic amino acid catabolism-related gene expression during CR but expression of lipid metabolism-related genes under AL diet. Together, our findings reveal new roles of adiponectin in the adaptive response to CR and highlight sex and diet as key determinants of adiponectin’s systemic and tissue-specific metabolic effects (Figure 6).

**Figure 6.**
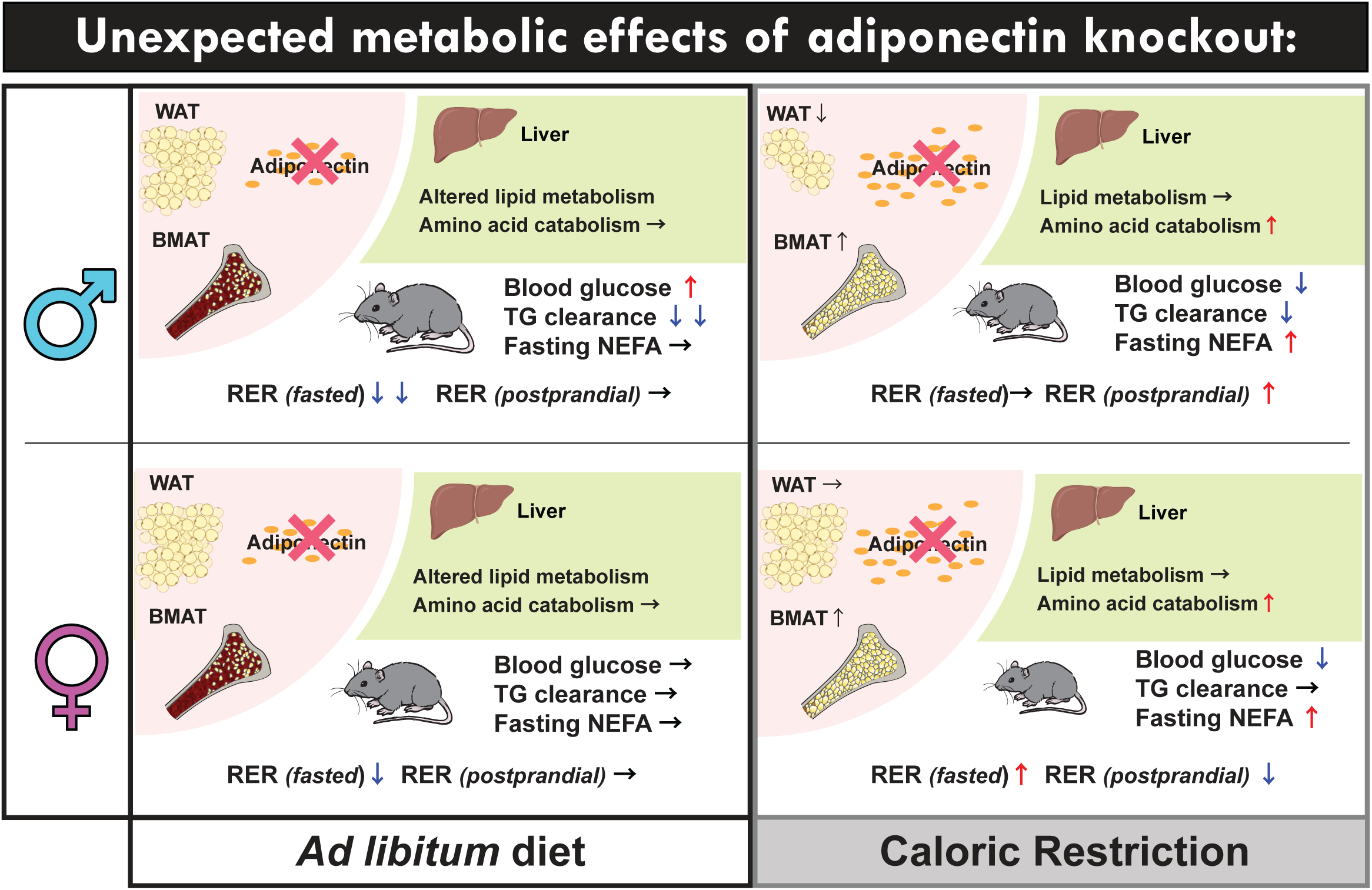
Adiponectin has distinct effects on lipid, amino acid, and glucose metabolism under AL and CR. Circulating adiponectin is secreted by WAT and BMAT. During CR, BMAT accumulates and contributes to increased plasma adiponectin, whereas WAT mass decreases in a sex-specific manner. Under AL diet, adiponectin KO alters hepatic lipid metabolism in both sexes, including effects on FA and sterol biosynthesis. Adiponectin KO also decreases RER during fasting (daytime) in AL males and females, suggesting increased lipid vs carbohydrate utilisation. However, some KO effects are sexually dimorphic: in AL males, KO causes mild hyperglycaemia, and impaired TG clearance, but these KO effects are absent in AL females. Under CR, adiponectin KO enhances hepatic amino acid catabolism through the regulation of gene expression. In contrast to AL, adiponectin KO promotes mild hypoglycaemia and increased fasting NEFA under CR and this may be driven, in part, by differential effects on RER in the fasted and postprandial states. Liver and WAT illustrations are from BioRender.com.

It is particularly striking that adiponectin KO lowers blood glucose levels in OGTT during CR, because adiponectin is typically regarded as an insulin-sensitising hormone that promotes glucose tolerance and prevents hyperglycaemia. However, on a normal, non-obesogenic diet, adiponectin KO mice have unaltered glucose tolerance, whether they are young, aged, or insulin-deficient^13,16,27–29^. Moreover, one human study found that CR *decreases* high-molecular weight adiponectin and that this is associated with increased glucose tolerance^26^. Thus, the impact of adiponectin deficiency on glucose homeostasis appears critically dependent on dietary context.

We show that, during CR, adiponectin KO lowers blood glucose without augmenting insulin secretion or insulin sensitivity, or impairing gluconeogenesis from glycerol. Instead, KO alters the changes in RER during transitions between fed and fasted states. In CR males, the rapid postprandial increase in RER is augmented in KO vs WT mice. This suggests that, during CR, adiponectin KO enhances carbohydrate utilisation after feeding, an effect that likely also occurs after oral glucose administration. This may explain why, in CR males, KO decreases fasting and peak glucose during the OGTT. In contrast, in CR females, adiponectin KO *attenuates* the postprandial rise in RER, which may explain why KO does not suppress OGTT peak glucose in CR females. Instead, KO suppresses the decline in RER as CR females progress from the fed (daytime) to fasted (nighttime) state. This suggests that, under CR, KO females resist the increase in lipid oxidation and/or sustain elevated carbohydrate oxidation during the onset of fasting. These effects may underlie the lower OGTT blood glucose and increased fasting NEFA observed in KO vs WT females under CR.

It remains less clear how KO increases fasting NEFA in CR males. This is unlikely to reflect greater lipolysis, because adiponectin KO does not alter adiposity or CR-induced fat loss. Future studies using labelled FA tracers would be helpful to determine how adiponectin KO impacts systemic and tissue-specific FA metabolism during CR.

We show that adiponectin KO enhances expression of genes involved in amino acid catabolism in both sexes, underscoring the robustness of this effect. This data suggests that KO males and females have increased reliance on amino acid catabolism to support gluconeogenesis, exemplified by increased expression of *Got1* and *Sds* in KO vs WT males during CR (Figure 5B, 5F, Supplemental Figure S4A, S6A-B). These KO effects are notable because gluconeogenesis from amino acids is less metabolically efficient than from glycerol^30^. Moreover, when lipid utilisation is constrained, amino acid metabolism becomes rate-limiting for gluconeogenesis, resulting in fasting hypoglycaemia^31^. We speculate that these hepatic effects also contribute to KO-induced hypoglycaemia and lower OGTT glucose during CR. Importantly, the RER for amino acid oxidation is ∼0.8 whereas for lipids it is 0.7; thus, if KO does cause a shift from lipid or glucose to protein oxidation during CR, this might not be detectable through systemic RER measurements. Further investigation will be required to determine the mechanism by which KO mice exhibit hypoglycaemia under CR.

The underlying mechanism whereby adiponectin KO enhances expression of genes involved in amino acid catabolism in both sexes remains unclear, with few reports linking adiponectin to amino acid catabolism. Peroxisome proliferator-activated receptor alpha (PPARα), a nuclear receptor activated by fatty acids, fasting, and CR, is critical for the hepatic response to fasting and regulates both lipid and amino acid metabolism^32–36^. Notably, adiponectin can also activate PPARα^37^. Consistent with this, we show that there is extensive overlap between PPARα target genes related to amino acid metabolism and the genes enriched in KO livers under CR (Supplementary Figure 6A and B; *Acmsd, Agxt2, Arg1, Cbs, Got1, Got2, Gls, Gls2, Hpd, Oat, Otc, Pah, and Tat)*^38^. Thus, one possibility is that adiponectin acts via PPARα to regulate hepatic amino acid metabolism under CR. Given that plasma adiponectin is inversely associated with circulating amino acid concentrations in humans^39^, future studies should further establish how adiponectin influences hepatic amino acid catabolism and whole-body amino acid metabolism during CR.

Our findings shed further light on other hepatic effects of CR and adiponectin. The adiponectin receptors have ceramidase activity, and many studies report that adiponectin suppresses hepatic ceramide content^40–43^. Yet we show no effect of adiponectin KO on hepatic ceramides, DHCs, and ceramide:DHC ratios in AL or CR mice, whether for total concentrations or for the individual species. This echoes previous studies finding limited or no effect of adiponectin on ceramides in SkM or the pancreas^13,29^. It may be that adiponectin’s hepatic ceramide-lowering actions are greatest in states of obesity, insulin resistance, and/or hyperinsulinaemia. Whatever the case, our data robustly establish that adiponectin does not always influence hepatic sphingolipids and does not contribute to their modulation during CR.

Our study also highlights roles of adiponectin in systemic lipid metabolism. We show that global adiponectin KO significantly reduces lipid clearance in males and the KO effect is stronger in AL than in CR males. This adiponectin KO effect in male mice occurs in many other contexts, including in aged males fed chow or high-fat diet^16^; in type 1 diabetic males^29^; and in males with tissue-specific adiponectin deletion in WAT or the kidney^44,45^. This relates to the ability of adiponectin to stimulate lipoprotein lipase activity^46^, which decreases with adiponectin deficiency in mice and humans^45,47^. No previous studies have assessed lipid tolerance in KO females; however, transgenically increased adiponectin improves lipid clearance in mice of both sexes^16,48^. Together, the present and previous observations suggest that increased lipid clearance is among the most-robust metabolic effects of adiponectin in males, whereas in females this is more context-dependent.

While we have focussed on the liver as the key target of CR and adiponectin, a recent study highlights roles for adiponectin in renal gluconeogensis^44^. This is notable because renal gluconeogenesis contributes to ∼40% of glucose production in starvation^49^ and can compensate for impaired hepatic gluconeogenesis^50,51^. Adiponectin also targets many other cells and tissues, including SkM, beta cells, the heart, endothelia, leukocytes, and adipose tissue. Thus, investigating the kidney and other targets may further resolve the mechanisms underlying adiponectin’s unexpected effects in CR.

Finally, across our experiments we consistently find that the impact of adiponectin deficiency is diet- and sex-dependent. This suggests that adiponectin’s metabolic actions depend critically on other variables influenced by these factors, including insulin, glucagon, leptin, ghrelin, glucocorticoids, oestrogens, and androgens. Indeed, each of these hormones exerts fundamental metabolic effects, including during CR^3^, while sex steroids often underpin sex differences in metabolic function^52^. Importantly, most preclinical studies of increased or decreased adiponectin action have been in male mice, overlooking potential sex differences. It will therefore be critical to further establish the extent of, and basis for, adiponectin’s diet-dependent, sexually dimorphic effects.

Together, our study demonstrates the role of adiponectin in tissue-specific and systemic metabolism differs significantly under AL and CR, providing insight into the potential evolutionary functions of this enigmatic hormone. Future research should seek to determine the molecular mechanisms by which adiponectin affects lipid, amino acid, and glucose metabolism under CR, and if adiponectin regulates other aspects of the CR response. Finally, while CR can serve as a paradigm to understand the fundamental roles of adiponectin, it is perhaps even more widely studied for its potential therapeutic benefits. In this light, it will be important to test if variation in adiponectin influences human responses to CR, time-restricted feeding, intermittent fasting, or other therapeutic nutritional interventions.

## 4. MATERIALS AND METHODS

Reagents and resources used in this study are described in Supplementary Table 5.

### 4.1. Animals

All mouse studies were approved by the University of Edinburgh Animal Welfare and Ethical Review Board and were conducted under project licenses granted by the UK Home Office. To generate *Adipoq* KO mice, sperm from transgenic *Adipoq*^tm1a^(KOMP)^Wtsi^ mice on a C57BL/6N background was purchased from The Knockout Mouse Project (KOMP) Repository (mouse colony TCPA0796, Toronto Centre for Phenogenomics, Toronto, Canada) and used by the Central Transgenic Core facility (University of Edinburgh) for *in vitro* fertilisation (IVF) of eggs from WT C57BL/6NCrl females. During IVF, Tat-Cre recombinase was injected into eggs for recombination at the LoxP sites^53^.. This excised *Adipoq* exon 3 at the single-cell stage, converting the *Adipoq*^tm1a^(KOMP)^Wtsi^ allele to the full KO allele. *Adipoq*^tm1b^(KOMP)^Wtsi^ heterozygotes were mated to generate KO and control (WT) mice. Littermate controls were used throughout, except for four separate C57BL/6NCrl males (which were included in the OGTT and ITT studies). Genotyping was done by Transnetyx (Cordova, Tennessee, USA).

Mice were housed on a 12 h light/dark cycle in a specific-pathogen-free facility with free access to water. Mice had free access to food unless they were undergoing CR, as described below. Sample sizes were determined by power calculations using G*Power software, with effect sizes based on previously published data for glucose tolerance and fat loss during CR^54^. Single mice were used as experimental units because mice were single housed. Comparisons were made between AL vs CR mice (within each genotype), and WT vs KO mice (within each diet). The exact number of mice is stated in each figure legend. Randomisation, blinding, and exclusion of mice from final analyses were as described^5^, with KO and WT mice randomly assigned to each diet. Seven mice were excluded because of confounding health and/or technical issues, including one KO AL male with oesophageal damage from OGTT gavage; four males (two WT AL, two WT CR) with gavage damage during OLTT; and two females (one KO, one WT) with unexplained excessive weight loss during CR. For some cohorts, specific tissues were not weighed at necropsy and therefore these mice have no data for these readouts; further details will be provided in the Source Data file.

### 4.2. Mouse CR studies

Mice were fed AL (Research Diets D12450B) or CR (Research Diets D10012703, administered at 70% of the average daily AL diet consumption) as described previously^5^. Mice were assessed non-invasively for body fat, lean mass, and free fluid weekly from weeks 0 to 4 of AL or CR feeding (∼9-13 weeks of age) using TD-NMR (Minispec LF90II; Bruker Optics, Billerica, MA, USA). Blood glucose was measured from weeks 0 to 4 using a OneTouch Verio Glucometer (LifeScan IP Holdings, Zug, Switzerland). Before necropsy, all mice were fasted for ∼6-12 h to ensure that any CR effects reflect chronic CR and not short-term fasting, as recommended^55^. At necropsy, mice were sacrificed via cervical dislocation and decapitation. Tissues were dissected, weighed, and stored in 10% formalin or on dry ice. Tissues in formalin were fixed at 4°C before being washed and stored in DPBS (Life Technologies). Tissues on dry ice were stored at -80°C prior to downstream analysis.

### 4.3. Adiponectin, NEFA, and TG analysis in plasma samples

Plasma concentrations of adiponectin (MRP300, Bio-Techne) or TG (Serum Triglyceride Determination Kit, TR0100, Sigma-Aldrich) were measured by ELISA or colourimetric assays as per manufacturer’s instructions. Plasma NEFA were measured as described^5^. Assay aborbances were measured using an Infinite 200 Pro plate reader (Tecan Life Sciences). Immunoblotting of plasma to confirm the presence or absence of adiponectin in WT and KO mice was done as described^54^. Total plasma protein was determined using total protein stain (926-11010, LiCor).

### 4.4. TG quantification in the liver

Approximately 100 mg of frozen liver tissue was placed in 1 mL cooled PBS containing 5% v/v IGEPAL CA-630 (Sigma, I8896) on ice. Tissue was homogenised using steel ball bearings and high-frequency shaking and this homogenate was centrifuged at 12,000 rcf for 10 minutes at 4°C to pellet debris. The supernatant was transferred to a new microtube and any floating lipid was resuspended. Pre-prepared 500 mg/dL TG standard (Sigma, 17811-1AMP) was serially diluted to 250, 125, 62.5, 31.25, 15.625 and 7.8125 mg/dL; a solution containing no TG standard was included as a negative control. TG concentrations in standards or liver homogenates (2 µL) were determined using an enzymatic colourimetric assay (Sentinel 17624H) according to the manufacturer’s instructions. Concentrations were normalised to the mass of liver used for each sample.

### 4.5. Metabolic tolerance tests

Mice were fed a half portion of food the evening (18:00) prior to the test. At 12:00 on the day of the test, basal blood glucose from lateral tail vein blood was measured using a OneTouch Verio glucometer; for OGTT and OLTT, a basal blood sample was also collected into EDTA-coated tubes (Microvette 16.444, Sarstedt, Leicester, UK). For OGTT, D-glucose (25% w/v in water) was then administered at 2 mg/g body mass by oral gavage. For OLTT, Intralipid was administered by gavage at 15 µL/g body mass. For ITT, Humulin (Eli Lilly and Company Ltd, Basingstoke, UK) was injected intraperitoneally at 0.7 mlU/g of lean mass (assessed by TD-NMR the day before ITT). For glycerol tolerance tests, glycerol (10% w/v in saline) was injected intraperitoneally at 1.5 mg/g body mass. Blood glucose was measured at the following timepoints post-substance administration: 15-, 30-, 60- and 120-minutes for OGTT and glycerol tolerance tests; and 15-, 30-, 60-, and 90-minutes for ITT. For OGTT and OLTT, tail vein blood was sampled at each timepoint in EDTA tubes and kept on ice until plasma isolation.

Plasma insulin during OGTT was measured using the Ultra-Sensitive Mouse Insulin ELISA Kit (#90080, ChrystalChem, Chicago, USA,) as per manufacturer’s instructions. HOMA-IR was calculated as described^56^.

### 4.6. Indirect calorimetry

Mice were housed individually in Promethion CORE System cages (Sable Systems International) from weeks 2-3 after beginning AL or CR feeding. Mice entered cages around 11:00 on day 1 and were housed for three nights (four days in total). Data are from the 48 h between 07:00 on day 2 and 07:00 on day 4 to allow environmental habituation during day 1. Before and after Promethion housing, mice were weighed and body composition determined by TD-NMR. Energy expenditure, O_2_ consumption, and CO_2_ production were analysed using ExpeData software as per manufacturer’s instructions.

### 4.7. Liver ceramide quantification

Liver ceramide and DHC content was determined using LC-MS (Lipidomics Core Facility; University of the Highlands and the Islands, Inverness, UK) as described previously^5^. In the final analysis, 26:0 ceramide values were excluded from three mice (two AL WT males and one CR KO male) because they were extreme statistical outliers (as identified using the Rout method with Q=1%, based on variance of combined values from across all diet and genotype groups). The ceramide:DHC ratio was also excluded for these mice.

### 4.8. Histology and assessment of myofibre cross-sectional area (CSA)

Formalin-fixed gastrocnemius muscles were paraffin-embedded by the Histology Core at The University of Edinburgh’s Shared University Research Facilities (SuRF). Sections (4 µm thick) were made at the widest lateral point of the muscle, de-waxed, stained with WGA-Alexa Fluor 488 (1:200) for 1 h, washed with PBS, stained with DAPI (1:100), and mounted with PermaFluor mounting medium. Three-to-seven images (10X objective) were taken per section using a Nikon Eclipse microscope and myofibre CSA analysed using ImageJ plug-in StarDist and MorphoJ. Particles >250 µm^2^ were used for calculation, with at least 689 fibres assessed per section.

### 4.9. RNA sequencing

For liver samples, RNA isolation, quantification, determination of the RNA Integrity Number (RIN), library preparation and sequencing were done as described^5^; data for livers of WT mice were also reported in this previous publication^5^.

Calculations were done using Eddie Mark 3, the University of Edinburgh’s high-performance computing cluster. After trimming by TrimGalore and quality control by FastQC, sequences were aligned to the mouse genome (mm10) with annotation data from the website of University of California Santa Cruz (https://hgdownload.soe.ucsc.edu) using STAR (v2.7.10a)^57^. Mapped reads were counted using featureCounts in Subread package (v1.5.2)^58^ and subsequent analyses performed using R, RStudio, GSEA, Cytoscape, and Metascape. Count data were normalised and analysed using DESeq2 (v1.44.0) to detect differentially expressed genes^59^.

PCA was done using the prcomp function for 500 genes, with the highest variation among samples after transforming the raw count data using the vst function from the DESeq2 package. The stats and ggplot2 packages were used for gene clustering and heatmapping. GO analysis was done with Metascape (https://metascape.org/gp/index.html#/main/step1)^60^. Metascape shows bar graph for viewing top non-redundant enrichment clusters, one per cluster. Each result shows Top 20 GO terms with –log 10 (p) > 2.0. For GESA, 1,000 gene-set permutations were used to estimate the null distribution for the data given the sample size, and “gene_set” was used for permutation type. But default setting was used otherwise.

### 4.10. Graphs and Statistical analysis

Data are shown as mean +/- SEM or as box-and-whisker plots. For the latter, the centre line is the median, the box extends from 25^th^ to 75^th^ percentile, the whiskers denote the minimum and maximum values, and individual data points are overlaid. Normal distributions were assumed for statistical analyses. Differences between experimental variables, and interactions between variables, were assessed by 3-way ANOVA (for data with three independent variables), 2-way ANOVA (for data with two independent variables), or mixed models (for repeated-measures analyses in which some data points were missing or had to be excluded for some mice). For RNA-seq data, a hypergeometric test was used to determine if the genes differentially expressed between KO and WT mice showed significant overlap between diet-sex subgroups. Further details are provided within each figure legend. A *P*-value <0.05 (after adjustment for multiple comparisons) was considered statistically significant. *P*-values are either shown directly or are indicated by * (p<0.05), ** (p<0.01), or *** (p<0.001).

### 4.11. Data availability

All source data from which the figures are based will be made available through University of Edinburgh DataShare. Raw data from RNA sequencing of livers of WT mice has been deposited at NCBI’s GEO repository with accession number GSE230402. Raw RNA sequencing data for KO livers will be deposited at NCBI’s GEO repository. Any other relevant data are available from the authors upon request.

## AUTHOR CONTRIBUTIONS

Contributions are based on the CRediT (Contributor Roles Taxonomy) and are as follows: ***Conceptualisation***, Y.M.I., K.C.C., R.J.S. and W.P.C.; ***Data curation,*** Y.M.I., K.C.C., R.J.S., and W.P.C.; ***Formal Analysis***, Y.M.I., K.C.C., R.J.S., M.B., H.K., P.D.W., and W.P.C.; ***Funding Acquisition***, Y.M.I., K.C.C., K.T., A.H.B., N.M.M., R.K.S. and W.P.C.; ***Investigation***, Y.M.I., K.C.C., R.J.S., D.M., E.J.B., S.A.F.X.Z, K.J.S., B.J.T., A.L., H.K., P.D.W., and W.P.C.; ***Methodology***, Y.M.I., K.C.C., R.J.S., H.K., P.D.W., N.M.M. and W.P.C.; ***Project administration,*** N.M.M., R.K.S. and W.P.C.; ***Resources***, N.M.M., R.K.S. and W.P.C.; ***Supervision,*** Y.M.I., K.J.S., N.M.M., R.K.S. and W.P.C.; ***Visualisation***, Y.M.I., K.C.C., R.J.S. and W.P.C.; ***Writing – Original Draft***, Y.M.I., K.C.C. and W.P.C.; ***Writing – Review & Editing***, Y.M.I., K.C.C, A.H.B., R.K.S. and W.P.C.

## Rights Retention Statement

For the purpose of open access, the author has applied a Creative Commons Attribution (CC-BY) licence to any Author Accepted Manuscript version arising from this submission.

## ACKNOWLEDGEMENTS

We gratefully acknowledge staff within Bioresearch & Veterinary Services (University of Edinburgh) for their expertise and assistance in mouse studies, including John Henderson, Vicky Walczak-Gillies, Anna Adamska, Will Mungall, Ami Onishi (supported by BHF grant RE/18/5/34216), Ailsa Travers, and other Central Transgenic Core staff.

## SUPPLEMENTARY INFORMATION

**Supplementary Figure S1.**
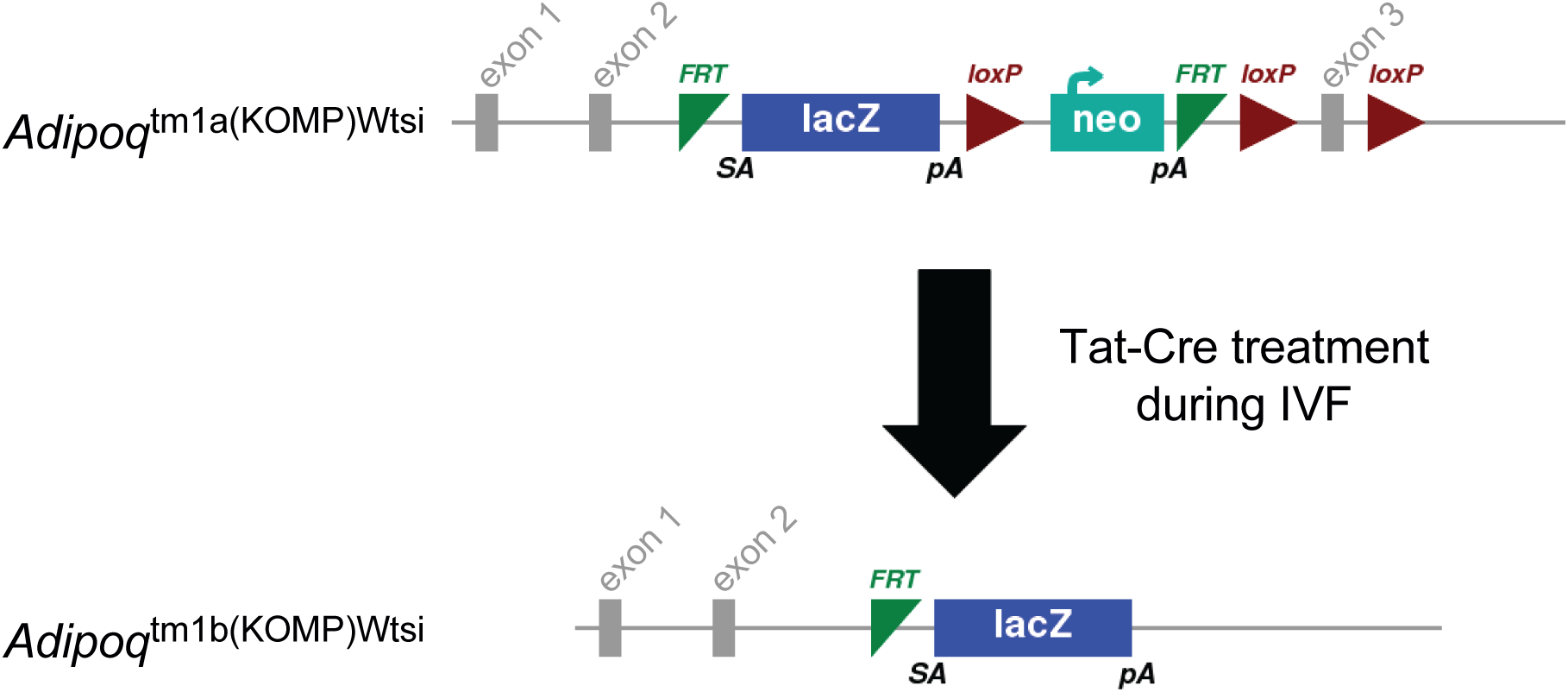
Adiponectin KO strategy. Sperm containing the *Adipoq*^tm1a^(KOMP)^Wtsi^ allele (*tm1a*) was used to fertilise eggs from WT C57BL/6NCrl females. During the IVF process, recombinant Tat-Cre recombinase was also injected, causing conversion to the *Adipoq*^tm1a^(KOMP)^Wtsi^ allele (*tm1b*) by excision of exon 3 of the *Adipoq* gene. Grey boxes are exons (numbered above); *FRT*, flippase recognition target; *lacZ*, open reading frame for beta-galactosidase reporter gene; *loxP,* recombination site for Cre recombinase; *neo,* open reading frame for neomycin resistance gene. Adapted from www.komp.org.

**Supplementary Figure S2.**
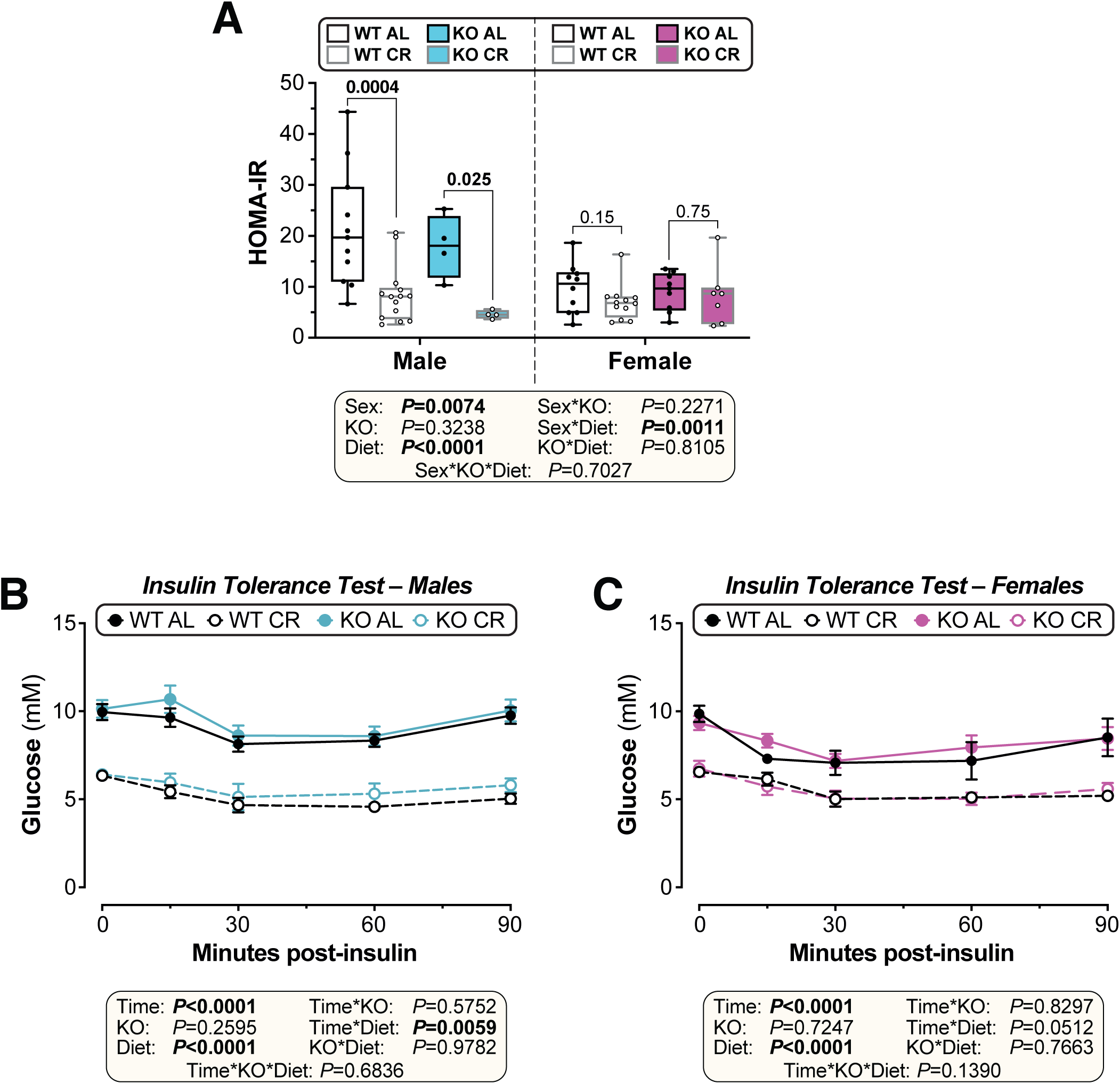
Adiponectin KO does not alter the effect of CR on insulin sensitivity. Male and female WT and *Adipoq* KO mice were fed AL or CR as described for Figure 1. **(A)** At 12.5 weeks of age (3.5 weeks of AL or CR diet) mice underwent an oral glucose tolerance test (OGTT). HOMA-IR of mice calculated from glucose and insulin concentrations during the OGTT. **(B-C)** At 12 weeks of age (3 weeks of AL or CR diet) mice underwent an insulin tolerance test (ITT). Blood glucose concentrations during the ITT are shown for males (B) and females (C). Data presentation and statistical analysis are as described for Figure 1, with data from the following numbers of mice per group: *male WT AL*, n= 11 (A) or 13 (B); *male WT CR,* n= 14 (A) or 13 (B); *male KO AL,* n= 4 (A) or 11 (B); *male KO CR,* n= 4 (A) or 10 (B); *female WT AL*, n= 10 (A) or 7 (C); *female WT CR,* n= 12 (A) or 11 (C); *female KO AL,* n= 9 (A) or 11 (C); *female KO CR,* n= 7 (A) or 10 (C).

**Supplementary Figure S3.**
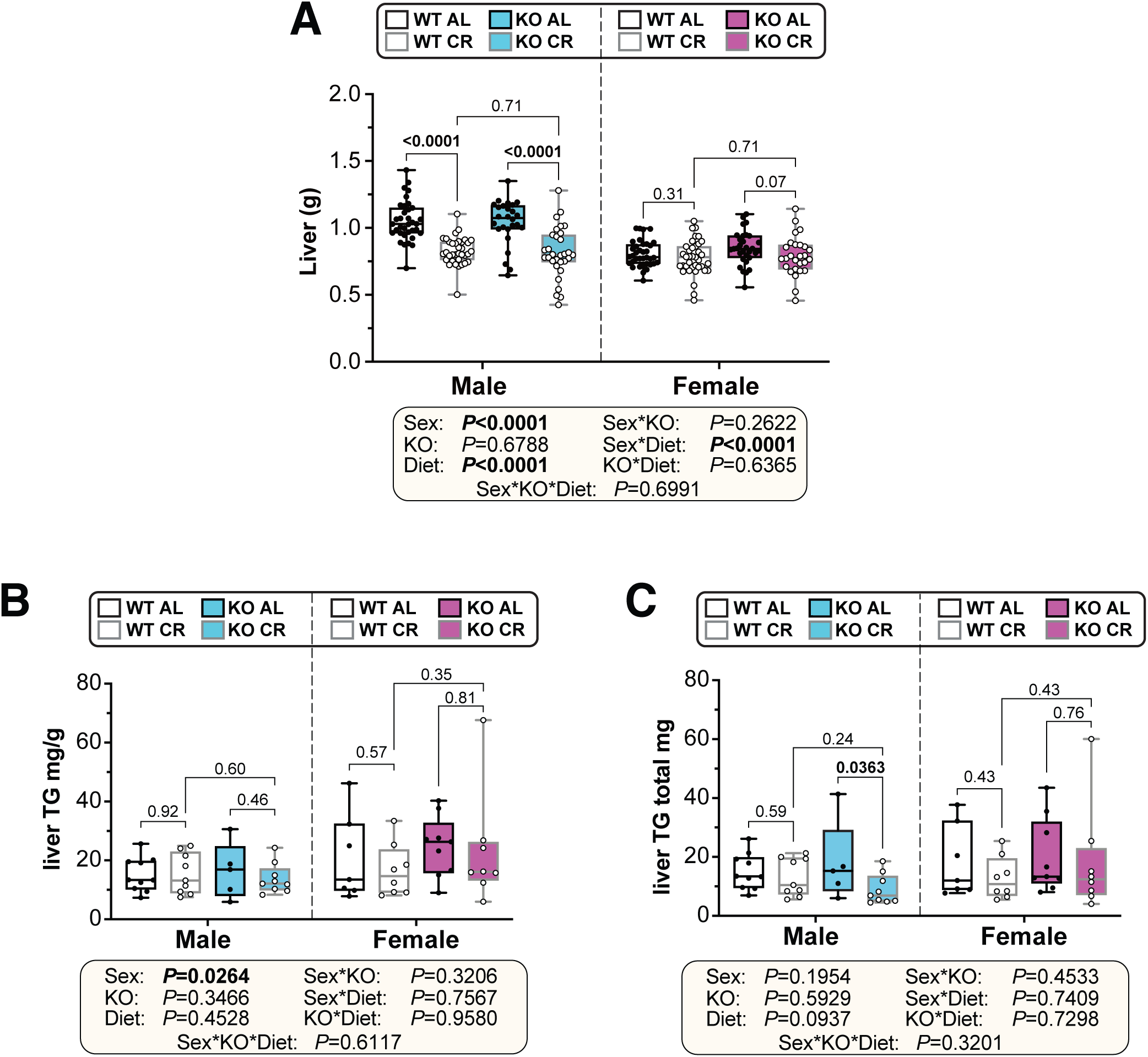
Adiponectin KO does not alter the effects of CR on liver mass or triglycerides. Male and female WT and *Adipoq* KO mice were fed AL or CR as described for Figure 1. At 13 weeks of age, mice were culled, and liver samples were collected. **(A)** Liver masses at necropsy, shown as box-and-whisker plots of the following numbers of mice per group: *male WT AL*, n=35 ; *male WT CR,* n=35; *male KO AL,* n=26; *male KO CR,* n=31; *female WT AL*, n=36; *female WT CR,* n=37; *female KO AL,* n=31; *female KO CR,* n=27. **(B, C)** TG content per unit weight of liver (B) and total hepatic TG content (C). Data are shown as box-and-whisker plots of the following numbers of mice per group: *male WT AL*, n=10; *male WT CR,* n=9; *male KO AL,* n=5; *male KO CR,* n=9; *female WT AL*, n=7; *female WT CR,* n=8; *female KO AL,* n=9; *female KO CR,* n=8. Statistical analyses are as described for Figures 1E-F.

**Supplementary Figure S4.**
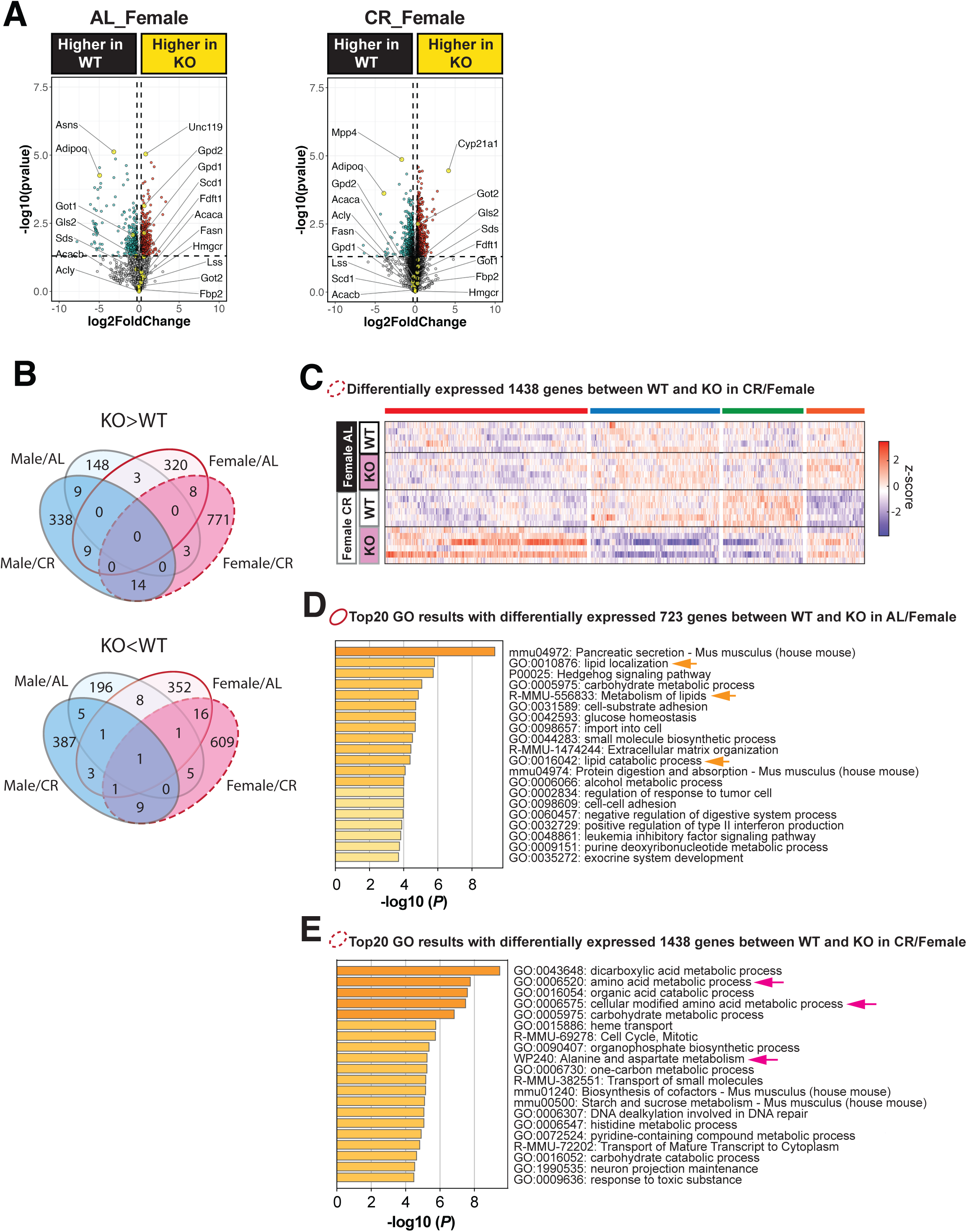
Effect of adiponectin KO on liver gene expression in females. Female WT and *Adipoq* KO mice were fed AL or CR and their livers analysed by bulk RNA-seq, as described for Figure 5. **(A)** Volcano plots showing differently expressed genes between WT and KO in female subgroups (AL Female, CR Female). Genes with rawp value < 0.05 and absolute fold change >1.2 are shown in blue or red dots. The name of some genes of interest are shown. **(B)** Overlap of genes differentially expressed between WT and KO in each subgroup (AL Male, CR Male, AL Female, and CR Female). Genes increased in KO vs WT (rawp<0.05, fold change >1.2) are shown in the upper Venn diagram and genes decreased in KO vs WT (rawp<0.05, fold change <-1.2) are shown in the lower Venn diagram. These Venn diagrams are the same to Figure 5C. **(C)** Clustered heat maps of the 1438 genes differentially expressed (rawp <0.05, absolute fold change >1.2) between WT and KO under CR in females, the gens included in the dashed red circles in the Venn diagrams in (B). **(D)** Results of the GO analyses conducted with the 723 genes differentially expressed between WT and KO in AL female, the genes included in the red circles in the Venn diagrams in (B). The top 20 clusters of GO results are shown. The yellow arrows highlight the GO terms related to lipid metabolism. **(E)** Results of the GO analyses conducted with the 1438 genes shown in (B) and (C). The top 20 clusters of GO results are shown. Pink arrows highlight GO terms related to amino acid metabolism. Sample numbers for the female liver RNA-seq are as follows: *Female WT AL*, n=5; *Female WT CR*, n=6; *Female KO AL*, n=6; *Female KO CR*, n=6.

**Supplementary Figure S5.**
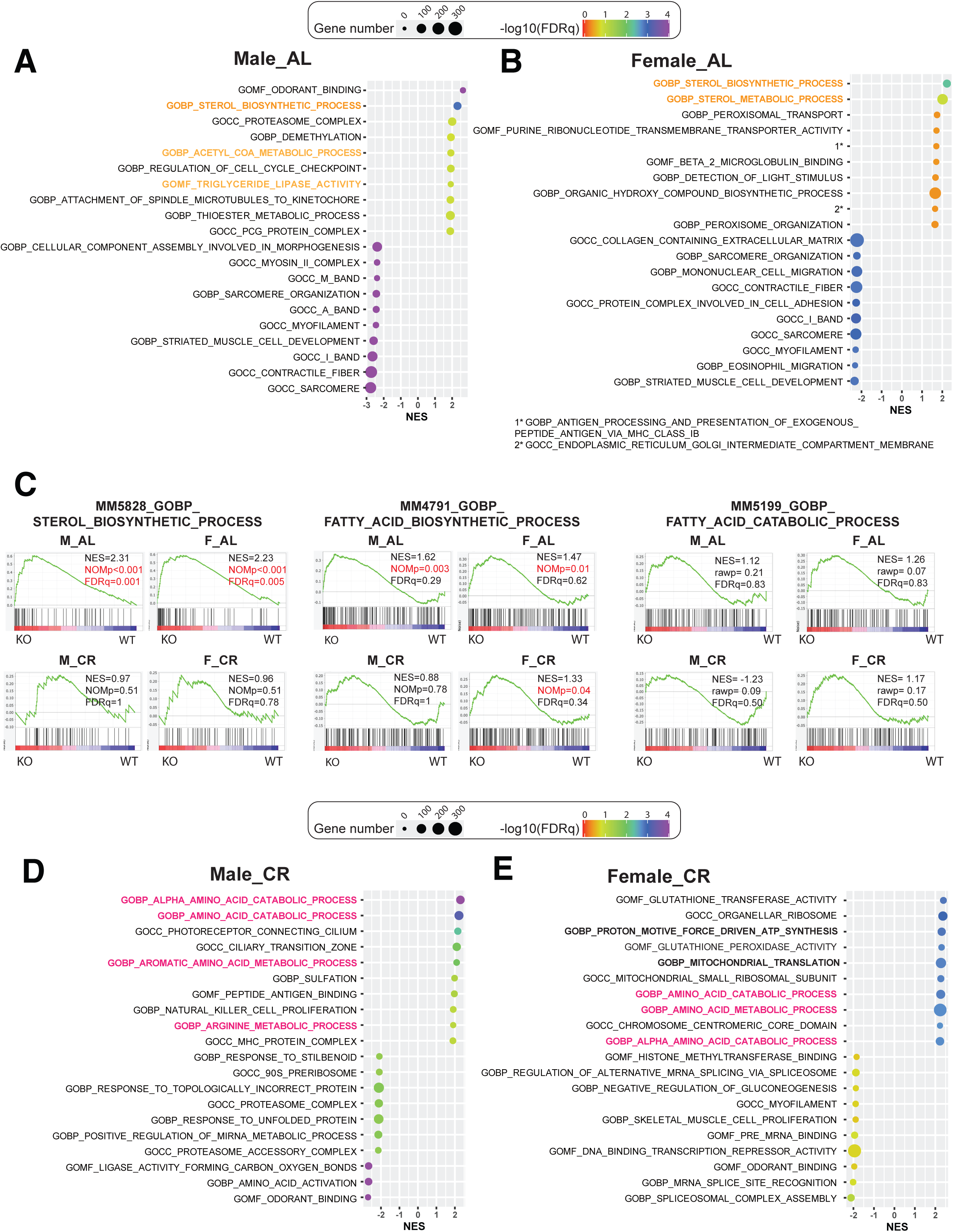
Lipid metabolism-related genes are major targets under AL while amino acid-related genes are major the targets under CR for Adiponectin. Male and Female WT and *Adipoq* KO mice were fed AL or CR and their livers analysed by bulk RNA-seq, as described for Figure 5. **(A-B)** Bubble plots of the top 10 GO gene sets (with lowest FDRq values) enriched in KO (NES>0) or WT (NES<0) within AL Male (A) or AL Female (B) subgroups. The GO terms are sorted by the NES values. Gene set names related to sterol metabolism are highlighted in orange, lipid catabolism in yellow **(C)** GSEA result between KO vs WT in AL Male (upper left), AL Female (upper right), CR Male (lower left), or CR Female (lower right) subgroups for MM5828_GOBP_STEROL_BIOSYNTHETIC_PROCESS (left), MM4791_GOBP_FATTY_ACID_BIOSYNTHETIC_PROCESS (middle), and MM5199_GOBP_FATTY_ACID_CATABOLIC_PROCESS (right). **(D-E)** Bubble plots of the top 10 GO gene sets (with lowest FDRq values) enriched in KO (NES>0) or WT (NES<0) within CR Male (D) or CR Female (E) subgroups. The GO terms are sorted by the NES values. Gene set names related to amino acid metabolism are highlighted in pink. For (A-E), data were extracted from GSEA results conducted with the M5 gene set library (m5.all.v2023.2.Mm.symbols.gmt). For (A-B) and (D-E), bubble size and colour coding for FDRq values are shown above the bubble plots. In cases where the FDR-q value is 0, a value of 0.0001 has been used (-log10(FDRq) = 4) to allow visualisation.

**Supplementary Figure S6.**
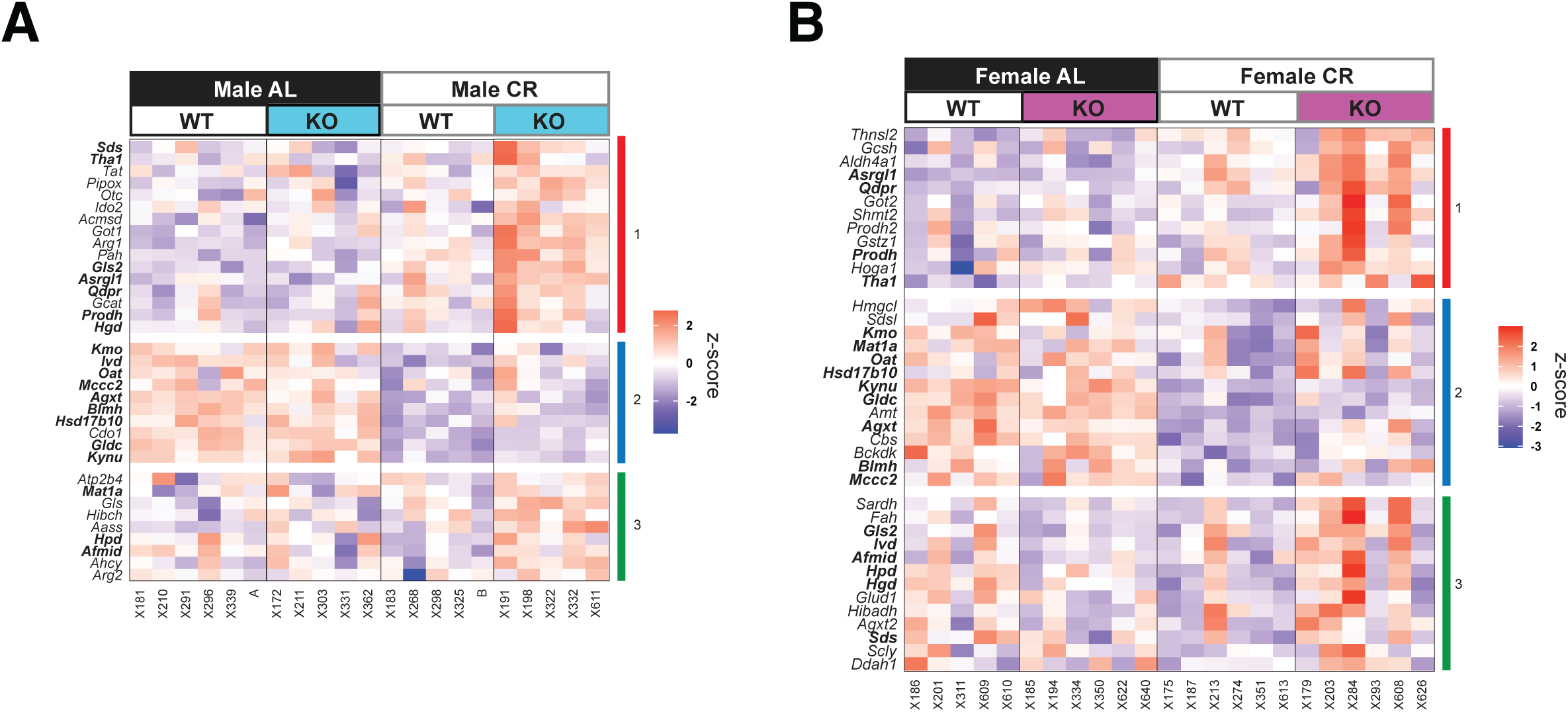
Adiponectin KO leads to upregulated expression of genes related to amino acid catabolism under CR. Male and Female WT and *Adipoq* KO mice were fed AL or CR and their livers analysed by bulk RNA-seq, as described for Figure 5. **(A, B)** Clustered heat map of the genes that showed core enrichment in MM10709_GOBP_ALPHA_AMINO_ACID_CATABOLIC_PROCESS gene set in comparisons between KO vs WT for CR Male (A) or CR Female (B) subgroups. The common genes between (A) and (B) are shown in bold. Sample numbers for liver RNA-seq are as follows: *Male WT AL*, n=6; *Male WT CR,* n=5; *Male KO AL*, n=5; *Male KO CR*, n=5; *Female WT AL*, n=5; *Female WT CR*, n=6; *Female KO AL*, n=6; *Female KO CR*, n=6.

**Supplementary Figure S7.**
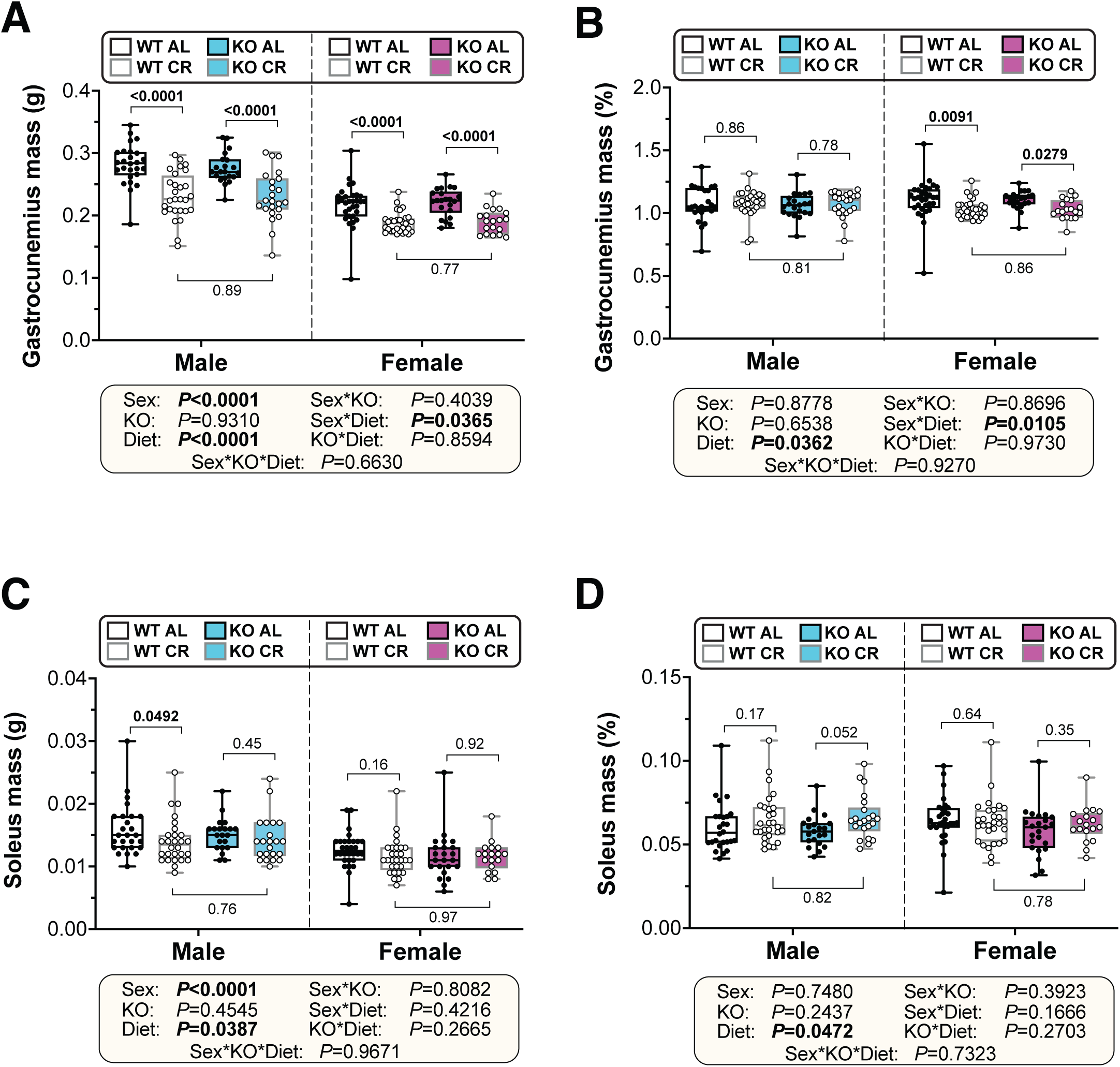
Adiponectin KO does not alter muscle mass under AL nor CR diet. Male and female WT and *Adipoq* KO mice were fed AL or CR as described for Figure 1. At 13 weeks of age, mice were culled, and SkM samples were collected. **(A-D)** Gastrocnemius and soleus masses were recorded at necropsy (13 weeks of age) and are shown as absolute mass (A and B) or % body mass (C and D). Data areown as box-and-whisker plots of the following numbers of mice per group *male WT AL*, n=26 ; *male WT CR,* n=28 ; *male KO AL,* n=21; *male KO CR,* n=23 for gastrocnemius or 22 for soleus; *female WT AL*, n=31 for gastrocnemius or 30 for soleus; *female WT CR,* n=30 for gastrocnemius or 29 for soleus; *female KO AL,* n=22 for gastrocnemius or 23 for soleus; *female KO CR,* n=19 for gastrocnemius or 18 for soleus. Statistical analyses were as described for Figures 1E-F.

**Supplementary Figure S8.**
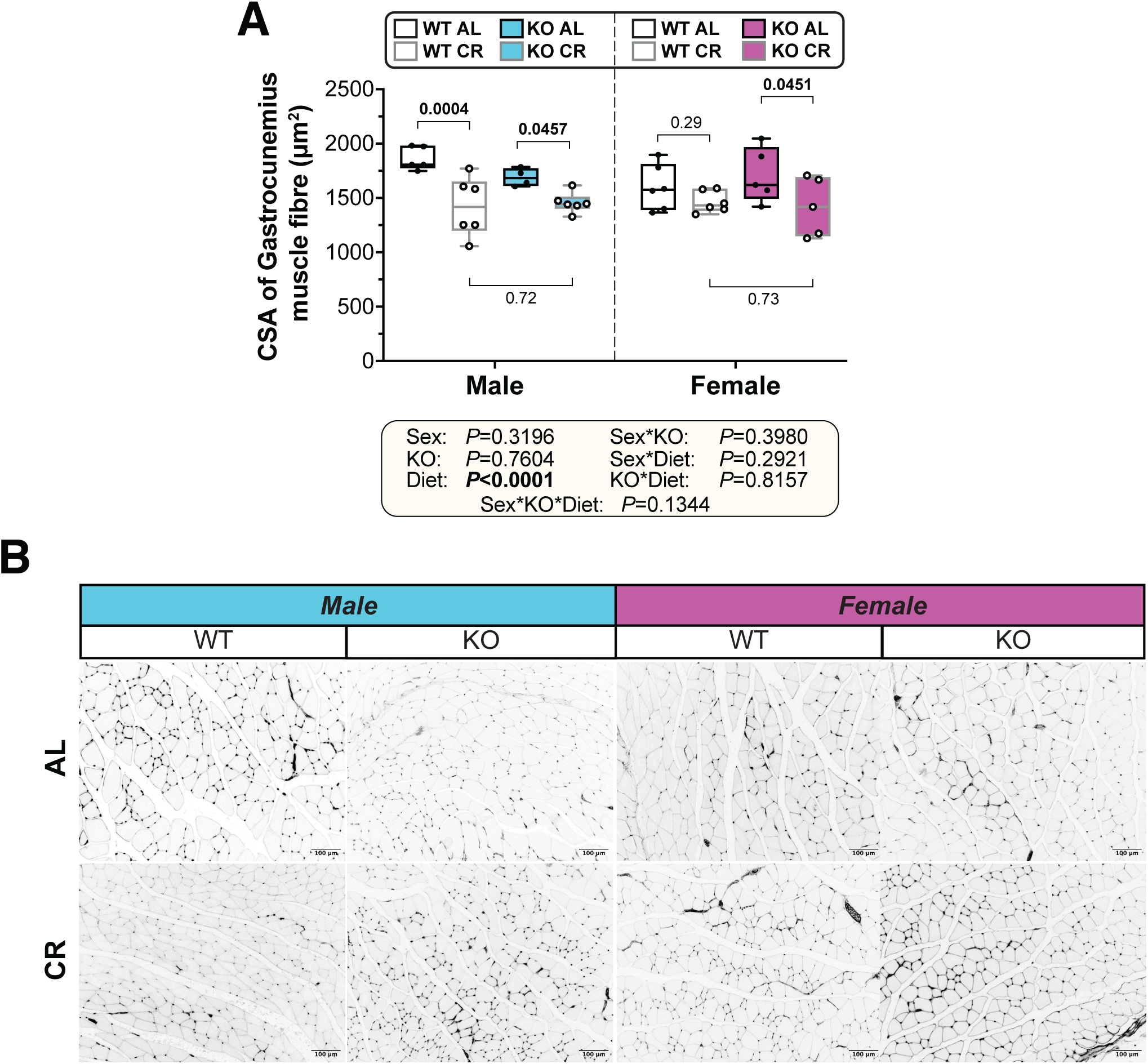
Adiponectin KO does not alter muscle fibre CSA in gastrocnemius under AL nor CR diet. Male and female WT and *Adipoq* KO mice were fed AL or CR as described for Figure 1. At 13 weeks of age, mice were culled, and SkM samples were collected. **(A)** CSA of gastrocnemius mass at the cull. **(B)** Representative images of gastrocnemius cross-sectional sections. Scale bars indicate 100um. Data are shown as box-and-whisker plots of the following numbers of mice per group *male WT AL*, n=5; *male WT CR,* n=6 ; *male KO AL,* n=4; *male KO CR,* n=6 ; *female WT AL*, n=6 ; *female WT CR,* n=6; *female KO AL,* n=5; *female KO CR,* n=5. Statistical analyses were as described for Figures 1E-F.

**Supplementary Figure S9.**
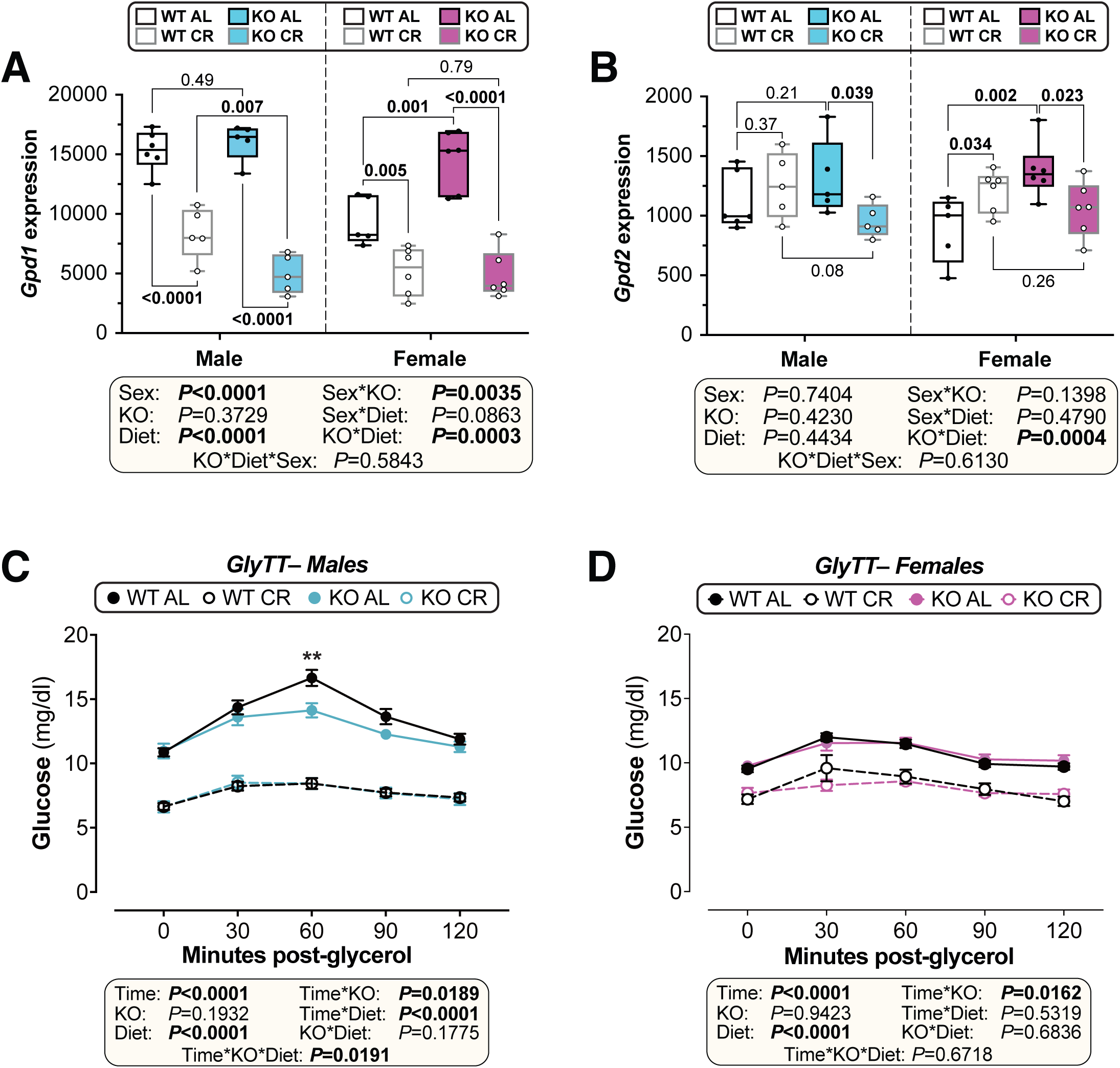
Effects of adiponectin KO on hepatic expression of gluconeogenesis-related genes and on glycerol-driven gluconeogenesis. Male and female WT and *Adipoq* KO mice were fed AL or CR and their livers analysed by bulk RNA-seq, as described for Figure 5. **(A-B)** Normalised count data for *Gpd1* (A) and *Gpd2* (B). Sample numbers for males and females are described in legends for Figure 5. Statistical analyses were as for Figures 1E-F. **(C-D)** Glycerol tolerance tests in male (C) and female (D) mice, shown as mean ± SEM of the following numbers of mice per group: *males*, n= 14 (WT AL, WT CR), 10 (KO AL), or 11 (KO CR); *females*, n= 16 (WT AL), 14 (WT CR), 10 (KO AL), or 7 (KO CR). Significant effects of diet, time, and/or genotype, and interactions thereof, were determined by 3-way ANOVA. Within each diet and sex, significant genotype effects at each time point were determined by 2-way ANOVA with Šidák’s multiple comparisons test; *\*\** (*P*<0.01).

### SUPPLEMENTARY TABLES

**Supplementary Table 1.**
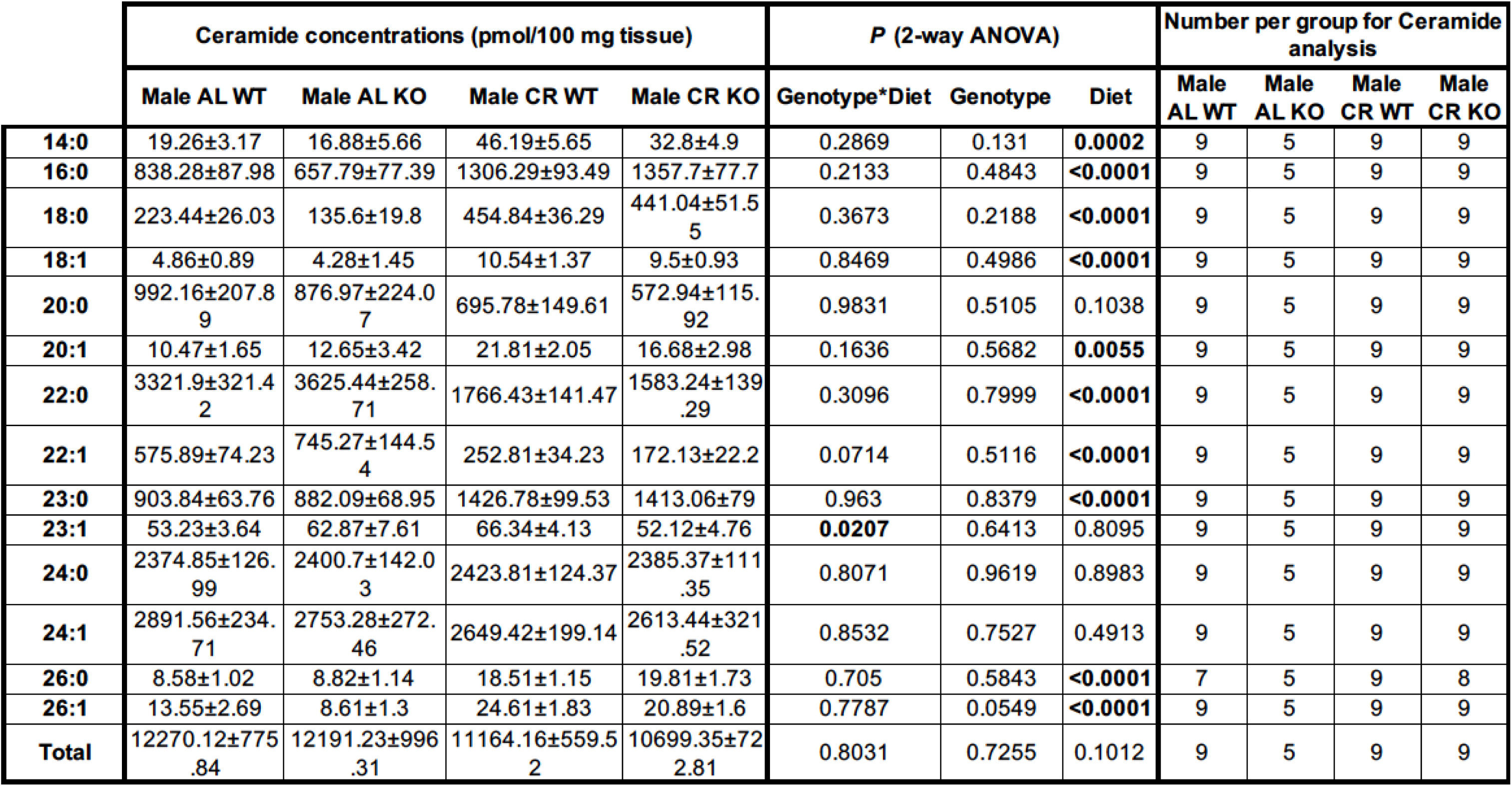
Ceramide concentrations for different sphingolipid species. The table shows LC-MS-measured concentrations of total and each species of ceramide. Average ± SEM are shown. Outlier identification analysis was conducted with the Rout method (Q=1%) and three outliers were excluded from the data for Ceramide 26:0.

**Supplementary Table 2.** DHC concentrations for different sphingolipid species. The table shows LC-MS-measured concentrations of total and each species of DHC. Average ± SEM are shown. DHC 26:0 was not detected for 7 samples and DHC 26:1 was not detected for 1 sample. The cells for those values were left blank.

|  | DHC concentrations (pmol/100 mg tissue) |  |  |  | P (2-way ANOVA) |  |  | Number per group for DHC analysis |  |  |  |
| --- | --- | --- | --- | --- | --- | --- | --- | --- | --- | --- | --- |
|  | Male AL WT | Male AL KO | Male CR WT | Male CR KO | Genotype*Diet | Genotype | Diet | Male AL WT | Male AL KO | Male CR WT | Male CR KO |
| **14:0** | 0.64±0.22 | 0.51±0.32 | 2.87±1.04 | 3.4±1.13 | 0.7226 | 0.83 | **0.0091** | 9 | 5 | 9 | 9 |
| **16:0** | 60.41±7.26 | 47.3±7.26 | 122.53±16.08 | 132.28±16.45 | 0.4311 | 0.9074 | **<0.0001** | 9 | 5 | 9 | 9 |
| **18:0** | 30.97±7.7 | 18.24±3.64 | 80.45±10.94 | 93.79±23.18 | 0.414 | 0.9846 | **0.0004** | 9 | 5 | 9 | 9 |
| **18:1** | 1.74±0.47 | 1.19±0.24 | 5.06±1.05 | 4.94±0.9 | 0.8025 | 0.6973 | **0.0003** | 9 | 5 | 9 | 9 |
| **20:0** | 52.11±12.53 | 42.45±3.36 | 58.57±5.09 | 58.21±11.99 | 0.6651 | 0.6408 | 0.3051 | 9 | 5 | 9 | 9 |
| **20:1** | 1.46±0.61 | 1.61±0.54 | 2.93±0.5 | 3.61±1.11 | 0.7433 | 0.6113 | **0.0409** | 9 | 5 | 9 | 9 |
| **22:0** | 179.2±21.41 | 169.51±18.83 | 111.89±8.09 | 102.26±15.3 | 0.9989 | 0.5775 | **0.0005** | 9 | 5 | 9 | 9 |
| **22:1** | 46.55±7.42 | 56.23±7.47 | 29.98±3.35 | 24.11±4.5 | 0.1995 | 0.7502 | **0.0003** | 9 | 5 | 9 | 9 |
| **23:0** | 44.12±5.5 | 46.4±6.76 | 108.17±6.31 | 102.8±16.27 | 0.7284 | 0.8884 | **<0.0001** | 9 | 5 | 9 | 9 |
| **23:1** | ND | ND | ND | ND |  |  |  | 0 | 0 | 0 | 0 |
| **24:0** | 103.03±10.96 | 119.56±14.06 | 163.76±10.26 | 147.9±17.77 | 0.2669 | 0.9814 | **0.0042** | 9 | 5 | 9 | 9 |
| **24:1** | 199.66±13.52 | 189.28±12.94 | 246.29±29.73 | 213.46±24.67 | 0.6476 | 0.3815 | 0.1561 | 9 | 5 | 9 | 9 |
| **26:0** | 0.45±0.14 | 0.34±0.19 | 0.24±0.06 | 0.27±0.05 | 0.5747 | 0.7487 | 0.2459 | 7 | 2 | 7 | 9 |
| **26:1** | 0.48±0.12 | 0.95±0.13 | 1.73±0.39 | 1.35±0.35 | 0.224 | 0.8971 | **0.0208** | 9 | 4 | 9 | 9 |
| **Total** | 303.52±22.99 | 309.73±26.94 | 411.96±37.38 | 362.99±41.44 | 0.4536 | 0.5607 | **0.0342** | 9 | 5 | 9 | 9 |

**Supplementary Table 3.** Ceramide:DHC ratio for different sphingolipid species. The table shows Ceramide:DHC ratio based on LC-MS-measured concentrations of total and each species of ceramide (Supplementary Table 1) and DHC (Supplementary Table 2). Average ± SEM are shown. Outlier identification analysis was conducted with the Rout method (Q=1%) and three outliers were excluded from the data for Ceramide 26:0. The Ceramide:DHC ratio data for these three samples were also excluded. DHC 26:0 was not detected for 7 samples and DHC 26:1 was not detected for 1 sample, and the correnponding ceramide:DHC ratio values were left as blank.

|  | Ceramide:DHC ratio |  |  |  | P (2-way ANOVA) |  |  | Number per group for Ceramide:DHC ratio |  |  |  |
| --- | --- | --- | --- | --- | --- | --- | --- | --- | --- | --- | --- |
|  | Male AL WT | Male AL KO | Male CR WT | Male CR KO | Genotype*Diet | Genotype | Diet | Male AL WT | Male AL KO | Male CR WT | Male CR KO |
| **14:0** | 68.16±22.08 | 110.07±41.48 | 47.55±18.94 | 30.1±13.88 | 0.2037 | 0.5962 | **0.0358** | 9 | 5 | 9 | 9 |
| **16:0** | 14.25±0.93 | 14.15±0.5 | 11.48±0.99 | 11.32±1.2 | 0.9785 | 0.9051 | **0.0143** | 9 | 5 | 9 | 9 |
| **18:0** | 8.89±0.95 | 7.74±0.57 | 6.2±0.6 | 7.33±1.66 | 0.3443 | 0.9937 | 0.2019 | 9 | 5 | 9 | 9 |
| **18:1** | 4.87±2.24 | 4.2±1.13 | 2.31±0.2 | 2.32±0.35 | 0.807 | 0.8153 | 0.1183 | 9 | 5 | 9 | 9 |
| **20:0** | 22.58±3.95 | 19.83±3.6 | 12.18±2.42 | 14.74±4.76 | 0.5172 | 0.9803 | 0.0659 | 9 | 5 | 9 | 9 |
| **20:1** | 22.35±7.78 | 10.25±3.75 | 9.12±1.48 | 7.8±1.54 | 0.2773 | 0.1785 | 0.118 | 9 | 5 | 9 | 9 |
| **22:0** | 20.23±2.16 | 22.01±1.88 | 16.05±1.31 | 20.16±4.7 | 0.7163 | 0.3627 | 0.3507 | 9 | 5 | 9 | 9 |
| **22:1** | 13.76±1.63 | 12.92±1.1 | 8.94±1.21 | 11.05±3.64 | 0.5546 | 0.7991 | 0.1849 | 9 | 5 | 9 | 9 |
| **23:0** | 22.37±2.46 | 20.91±3.93 | 13.66±1.47 | 17.55±3.43 | 0.3622 | 0.676 | **0.0454** | 9 | 5 | 9 | 9 |
| **23:1** |  |  |  |  |  |  |  | 0 | 0 | 0 | 0 |
| **24:0** | 24.54±2.31 | 21.15±2.37 | 15.24±1.25 | 18.52±2.79 | 0.168 | 0.9813 | **0.0173** | 9 | 5 | 9 | 9 |
| **24:1** | 15.05±1.66 | 14.52±0.88 | 11.75±1.34 | 15.29±3.82 | 0.4364 | 0.5645 | 0.6274 | 9 | 5 | 9 | 9 |
| **26:0** | 49.13±12.55 | 31.53±16.3 | 93.4±14.95 | 97.45±18.84 | 0.6132 | 0.7512 | **0.0174** | 5 | 2 | 7 | 8 |
| **26:1** | 43.61±10.49 | 11.01±2.66 | 19.67±3.98 | 27.85±8.8 | **0.0304** | 0.1824 | 0.6935 | 9 | 4 | 9 | 9 |
| **Total** | 41.58±2.93 | 40.05±3.1 | 28.9±3.1 | 34.54±6.37 | 0.4412 | 0.6574 | 0.0574 | 9 | 5 | 9 | 9 |

**Supplementary Table 4.** Genes not shown in the volcano plots in Figure 5B. . The list of genes not shown in the volcano plots in Supplementary Figure 5B because the absolute log2FoldChange is over 10 or -log10(pvalue) is over 7.5. For CR/Females no genes meet these criteria (“n/a/”

| Diet/Sex | geneID | baseMean | log2 Fold-Change | lfcSE | stat | pvalue | padj |
| --- | --- | --- | --- | --- | --- | --- | --- |
| AL/Male | *3425401B19Rik* | 5.81 | -19.2 | 2.50 | -7.65 | 1.97E-14 | 3.38E-10 |
|  | *Glycam1* | 10.96 | -32.4 | 4.48 | -7.21 | 5.39E-13 | 4.61E-09 |
| CR/Male | *Myl2* | 9.19 | -19.7 | 2.65 | -7.43 | 1.11E-13 | 1.90E-09 |
|  | *Akp3* | 8.16 | 15.8 | 2.66 | 5.93 | 3.08E-09 | 2.63E-05 |
| AL/Female | *Glycam1* | 10.96 | -20.3 | 4.39 | -4.63 | 3.69E-06 | 0.052 |
| CR/Female | n/a | n/a | n/a | n/a | n/a | n/a | n/a |

**Supplementary Table 5.** Reagents and resources used in this study.

| REAGENT or RESOURCE | SOURCE | IDENTIFIER |
| --- | --- | --- |
| **CHEMICALS, KITS and ANTIBODIES** |  |  |
| D-Glucose | Sigma-Aldrich, Poole, UK | G8270 |
| Humulin | Eli Lilly and Company, Indiana, USA | HI0210 |
| IntraLipid | Sigma-Aldrich, Poole, UK | L141-100ML |
| Glycerol | Fisher Scientific, Loughborough, UK | G/P450/05 |
| Chloroform | Fisher Scientific, Loughborough, UK | C/4960/17 |
| TRIzol reagent | Life Technologies, Paisley, UK | 15596018 |
| Serum Triglyceride Determination Kit | Sigma-Aldrich, Poole, UK | TR0100 |
| Triglycerides TG enzymatic colorimetric Trinder assay | Sentinel Diagnostics, Milan, Italy | 17624H |
| Ultra Sensitive Mouse Insulin ELISA Kit | ChrystalChem, Chicago, IL, USA | 90080 |
| NEFA assay reagents | FUJIFILM Wako Chemicals Europe GmbH (Neuss, Germany) | 434-91795<br>436-91995<br>270-77000 |
| Qiagen RNeasy kit | Qiagen Ltd, Manchester, UK | 74106 |
| Microvette EDTA capillary tubes | Sarstedt, Leicester, UK | 16.444 |
| Anti-adiponectin (rabbit polyclonal Ab) | Sigma-Aldrich, Glasgow, UK | A6354 |
| **HISTOLOGICAL REAGENTS** |  |  |
| WGA-Alexa Fluor 488 | Thermo Fisher Scientific, Altrincham, UK | W11261 |
| DAPI | Thermo Fisher Scientific, Altrincham, UK | 62248 |
| PermaFlour mounting medium | Fisher Scientific, Loughborough, UK | 12695925 |
| **MOUSE MODELS** |  |  |
| Mouse: C57BL/6NCrl | Charles River, Edinburgh, UK | 027 |
| Adipoq KO | Described in "Animals" |  |
| **SOFTWARE & PROGRAMS** |  |  |
| Prism | GraphPad Software, LLC | V10.2.2 |
| Macro Interpreter | For analysis of Promethion data | V2.46 |
| Fiji/Image J | http://imagej.net | V2.14.0/1.54f |
| StarDist | https://imagej.net/plugins/stardist |  |
| MorphoJ | https://morphometrics.uk/MorphoJ_page.html |  |
| TrimGalore | https://www.bioinformatics.babraham.ac.uk/projects/trim_galore/ | V0.6.6 |
| FastQC | https://www.bioinformatics.babraham.ac.uk/projects/fastqc/ | v0.11.7 |
| STAR | Supplementary reference <sup>1</sup> | v2.7.10a |
| Subread | Supplementary reference <sup>2</sup> | v1.5.2 |
| R | https://www.r-project.org/ | v4.4.0 |
| RStudio | https://posit.co/download/rstudio-desktop/ | v2022.12.0+353 |
| DESeq2 | Supplementary reference <sup>3</sup> | v1.44.0 |
| ggplot2 |  | V3.5.1 |
| Metascape | Supplementary reference <sup>4</sup> | V3.5.20240101 |
| Cytoscape | Supplementary reference <sup>5</sup> | V3.10.2 |
| **OTHER** |  |  |
| Promethion CORE System | Sable Systems International (Las Vegas, USA) | ExpeData software v1.9.27 |
| OneTouch Verio Glucometer | LifeScan Inc., Zug, Switzerland | User's manual here |
| TD-NMR | Bruker Optics, Billerica, MA, USA | Minispec LF90II |
| Nikon microscope | Nikon, Tokyo, Japan | Nikon Eclipse Ti |
| CellCrusher | CellCrusher Limited, Schull, Ireland | WC Kit |

